# SAGA1 and SAGA2 localize the starch sheath to the pyrenoid in *Chlamydomonas reinhardtii*

**DOI:** 10.64898/2026.01.27.702031

**Authors:** Victoria L. Crans, Micah I. Burton, Aastha Garde, Lianyong Wang, Martin C. Jonikas

## Abstract

Most algae enhance their CO_2_ assimilation by concentrating CO_2_ within the pyrenoid, a biomolecular condensate that contains the CO_2_-fixing enzyme Rubisco. Many pyrenoids are surrounded by a starch sheath that is thought to slow the escape of CO_2_ from the pyrenoid, but how the starch sheath is localized to the pyrenoid remains poorly understood. Here, in the leading model alga *Chlamydomonas reinhardtii*, we find that the protein SAGA2 is necessary for early pyrenoid starch sheath biogenesis and works redundantly with its homolog, SAGA1, to localize the starch sheath to the pyrenoid. SAGA2 and SAGA1 were enriched in different regions of the pyrenoid-starch sheath interface, suggesting that they play complementary roles. Both *saga2* and *saga1* mutants showed defects in starch sheath coverage early during pyrenoid formation that were improved at a later timepoint. Strikingly, a *saga1;saga2* double mutant did not have a starch sheath around the pyrenoid and showed decreased overall starch content. SAGA1 and SAGA2 starch-binding domains bound to starch, the starch mimic molecule β-cyclodextrin, and the starch precursor molecule maltoheptaose, suggesting a role for SAGA1 and SAGA2 in starch granule initiation. We propose a model where SAGA1 and SAGA2 each locally prime starch sheath initiation in a distinct region of the pyrenoid surface by enriching starch precursor molecules around the pyrenoid. These findings advance the understanding of algal starch sheath biogenesis and provide insights into the associations between biomolecular condensates and other cellular structures.

**Significance Statement:** Eukaryotic algae enhance their carbon assimilation using an organelle called the pyrenoid, where concentrated CO_2_ is supplied to the CO_2_-fixing enzyme Rubisco. In many algae, a starch sheath surrounding the pyrenoid is thought to enhance CO_2_ fixation, but how starch is localized to pyrenoids is unknown. Here, we show that two proteins, SAGA1 and SAGA2, each bind to starch precursor molecules and redundantly localize starch to the pyrenoid in the alga *Chlamydomonas reinhardtii*. Our results suggest that SAGA1 and SAGA2 promote starch sheath initiation at the pyrenoid, rather than merely tethering starch, as previously thought. This work advances the understanding of the proteins and molecular mechanisms involved in pyrenoid starch sheath biogenesis and lays the foundations for their further study.

## Introduction

Approximately one-third of photosynthetic CO_2_ fixation occurs in the algal pyrenoid (1), an organelle that contains the CO_2_-assimilating enzyme Ribulose-1,5-bisphosphate carboxylase/oxygenase (Rubisco) (2, 3). There is interest in understanding how pyrenoids function because they mediate most of the CO_2_ assimilation in the oceans (4–8). In addition, engineering a pyrenoid into land plants is currently being pursued as a strategy for increasing crop yields (9–11).

Most of the molecular understanding of the pyrenoid comes from the leading model alga *Chlamydomonas reinhardtii* (Chlamydomonas hereafter). The Chlamydomonas pyrenoid is made up of three sub-compartments: the phase-separated (12–15) spheroidal Rubisco matrix (16); matrix-traversing membranes derived from the thylakoid sheets, which form tubules in Chlamydomonas (17, 18); and a surrounding starch sheath (19) (Fig. 1*A*). Each of these sub-compartments is thought to play a critical role in pyrenoid function: the Rubisco matrix is the site where CO_2_ is converted into sugar precursors that are used to generate biomass (20); the thylakoid tubules are thought to be the conduits that deliver concentrated CO_2_ to the Rubisco matrix (21–23); and the starch sheath is thought to improve the efficiency of CO_2_ concentration by acting as a barrier to slow the leakage of CO_2_ from the Rubisco matrix (23, 24).

**Figure 1.**
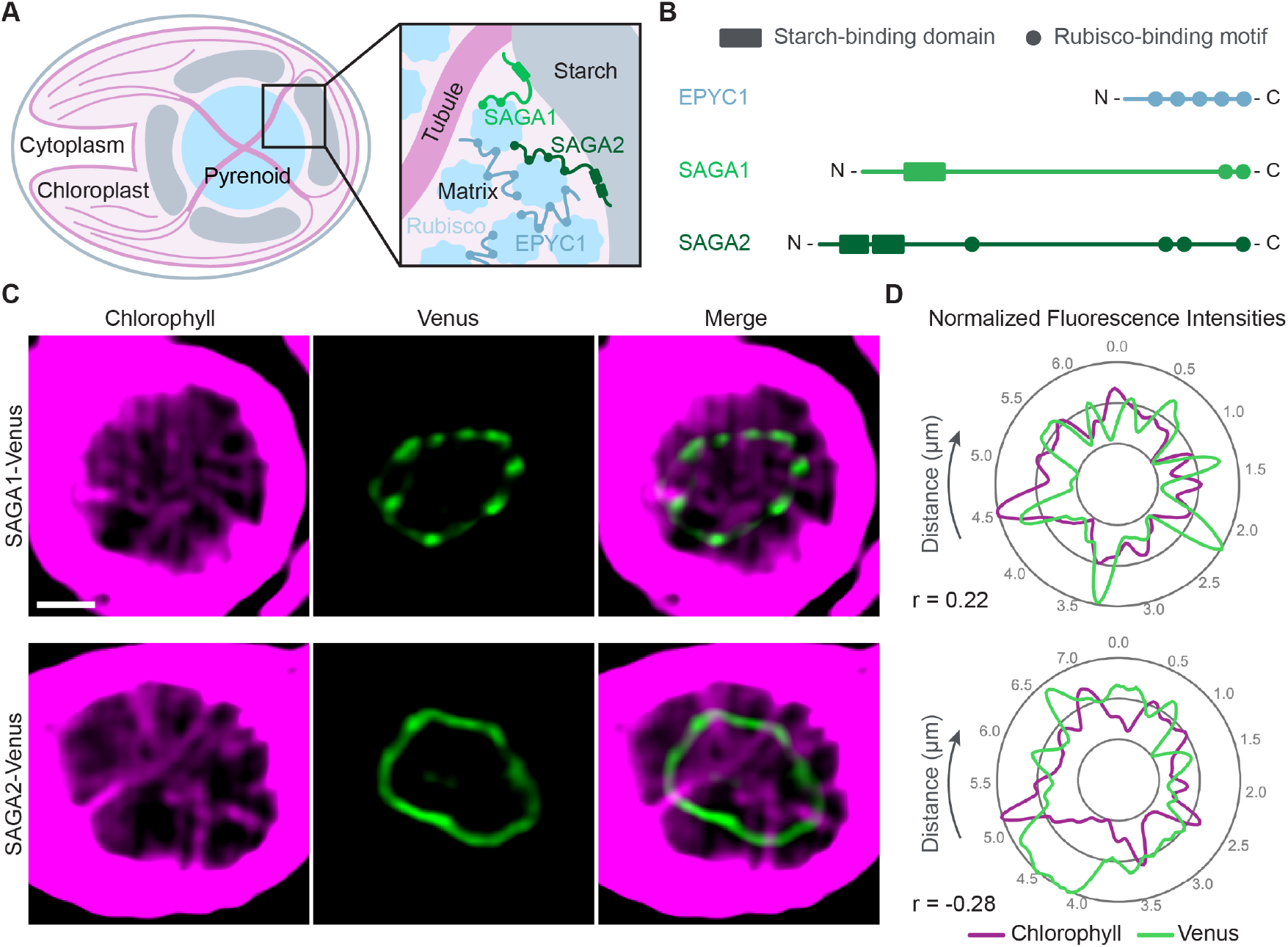
SAGA1 and SAGA2 localize to the matrix-starch interface of the pyrenoid. (A) Cartoon of a wild-type Chlamydomonas cell depicting the substructures of the pyrenoid and the localizations of SAGA1 and SAGA2. (B) The Rubisco linker protein EPYC1, SAGA1, and SAGA2 all contain copies of a Rubisco-binding motif that is required for pyrenoid formation. Additionally, SAGA1 and SAGA2 both contain predicted CBM20 starch-binding domains. (C) Representative confocal images of *saga1;SAGA1-Venus* and *saga2;SAGA2-Venus*. Scale bars = 1 µm. (D) Quantification of normalized fluorescence values for SAGA1-Venus, SAGA2-Venus, and chlorophyll autofluorescence. Graphs depict values for normalized fluorescence intensities around the periphery of the pyrenoids shown in C. Pearson’s correlation coefficients (r) for these measurements are labeled below the graphs. Images of additional pyrenoids and graphs of the corresponding normalized fluorescence values can be found in Fig. S1.

A role for starch as a CO_2_ leakage barrier is supported by in vitro, in vivo, and in silico data. Starch has been shown to have low permeability to gas diffusion in vitro (25, 26), and a Chlamydomonas mutant lacking a starch sheath cannot grow efficiently under limiting CO_2_ conditions (24). Additionally, modeling suggests that a CO_2_ diffusion barrier surrounding the matrix can increase the efficiency and efficacy of CO_2_ concentration within the pyrenoid (23).

If the starch sheath is indeed acting as a barrier to CO_2_ escape, it is likely that there would be cellular mechanisms dedicated to the proper localization of the starch sheath to the periphery of the Rubisco matrix. However, the mechanisms by which the starch sheath is localized to and shaped around the pyrenoid remain largely unknown.

Recent work identified two Chlamydomonas pyrenoid proteins that were proposed to localize the starch sheath to the periphery of the Rubisco matrix by physically connecting the two compartments (27). Meyer et al. (2020) discovered a Rubisco-binding motif that is both necessary and sufficient for targeting proteins to the pyrenoid (27) and is required for pyrenoid assembly (28). The motif is present on multiple pyrenoid proteins, including Essential PYrenoid Component 1 (EPYC1, Cre10.g436550), which binds Rubisco together to form the pyrenoid matrix (1, 28), and two proteins that localize to the starch-matrix interface of the pyrenoid: StArch Granules Abnormal 1 (SAGA1, Cre11.g467712 (29)) and its homolog, SAGA2 (Cre09.g394621 (27)) (Fig. 1 *A* and *B*). In addition to their Rubisco-binding motifs, SAGA1 and SAGA2 both have predicted carbohydrate-binding module (CBM)-20 starch-binding domains (27). The presence of the Rubisco-binding motifs and putative starch-binding domains on these two proteins, along with their localization to the starch-matrix interface, suggested that SAGA1 and SAGA2 could play roles in connecting the starch sheath to the Rubisco matrix.

In accordance with a potential role for SAGA1 in localizing the starch sheath to the Rubisco matrix, a *saga1* mutant from a Chlamydomonas insertional mutant library (30) was previously shown to have abnormal starch sheaths (29). This mutant had a severe growth defect at low levels of CO_2_, had multiple pyrenoid matrix condensates that lacked thylakoid tubules, and had thin and elongated starch plates around its pyrenoid condensates. Despite these morphological defects, the *saga1* mutant still had starch associated with its pyrenoid condensates, suggesting either that SAGA1 does not localize the starch sheath to the matrix or that it performs this function redundantly with one or more other proteins. SAGA2 is a promising candidate for this role, but a Chlamydomonas *saga2* mutant has not been previously characterized.

Recent work in *Arabidopsis thaliana* (Arabidopsis hereafter) supports the hypothesis that SAGA1 and SAGA2 function in localizing starch to the Rubisco matrix. Atkinson et al. (2024) found that the expression of SAGA1 and SAGA2 in Arabidopsis promoted the localization of starch within and around engineered Rubisco condensates in these plants (31). However, the specific role of SAGA2 and its functional relationship to SAGA1 remain unknown, motivating further study of these components in Chlamydomonas.

Here, we advance the understanding of the functions and relationship of SAGA1 and SAGA2 by characterizing the cellular localizations and loss-of-function phenotypes of these proteins in Chlamydomonas, and by investigating their ability to bind starch and starch precursors in vitro. Our results show that SAGA1 and SAGA2 act redundantly to localize starch granules to the pyrenoid matrix in Chlamydomonas and suggest that they localize starch by promoting the initiation of starch granules at the matrix surface. These findings further the understanding of the molecular interactions that are necessary to assemble a functional pyrenoid.

## Results

### SAGA2 is depleted from regions where membranes enter the pyrenoid matrix

SAGA1 and SAGA2 were both previously found to localize in ring-like patterns at the starch-matrix interface of the pyrenoid, with SAGA1 forming a punctate pattern (18, 27, 29) and SAGA2 having a more uniform distribution around the matrix periphery (27). Additionally, the SAGA1 puncta were shown to colocalize with chlorophyll autofluorescence around the edge of the pyrenoid, which corresponds to thylakoid tubule entry points, suggesting that SAGA1 localizes to gaps in the starch sheath where tubules enter the pyrenoid (18, 29).

To further investigate the localization pattern of SAGA2 in comparison to SAGA1, we used superresolution confocal microscopy to visualize SAGA1-Venus and SAGA2-Venus along with tubule chlorophyll autofluorescence (Fig. 1*C* and Fig. S1). By examining the spatial correlation between pixel intensities of Venus and chlorophyll fluorescence, we found that SAGA1-Venus on average had a positive correlation with tubule chlorophyll autofluorescence (Pearson correlation 0.19), while SAGA2-Venus on average had a negative correlation with tubule autofluorescence (Pearson correlation −0.27; Fig. 1*D* and Fig. S1). Our results indicate that whereas SAGA1 is enriched at the starch-tubule-matrix junctions of the pyrenoid, SAGA2 is depleted from these regions, validating previous observations and supporting the idea that SAGA1 and SAGA2 play complementary roles in starch sheath biogenesis.

### A *saga2* mutant does not have a detectable growth defect under standard photoautotrophic conditions

To investigate whether SAGA2 is necessary for starch sheath assembly in Chlamydomonas, we obtained a *saga2* mutant from the Chlamydomonas CLiP genome-wide mutant library (32). We verified the presence of an insertional cassette in the *SAGA2* gene in this mutant using PCR (Fig. S2 and Table S1). We then performed spot test assays to analyze the photoautotrophic growth of the *saga2* mutant under a photon flux of 100 μmol⋅m^−2^⋅s^−1^ at high (3% v/v), low (0.04% v/v) and very low (<0.004% v/v) levels of CO_2_ (Fig. 2*A* and Fig. S3). We found that the *saga2* mutant grew similarly to wild type under all growth conditions tested, suggesting that the mutant has a functional pyrenoid and that SAGA2 is not necessary for growth under limiting CO_2_ conditions.

**Figure 2.**
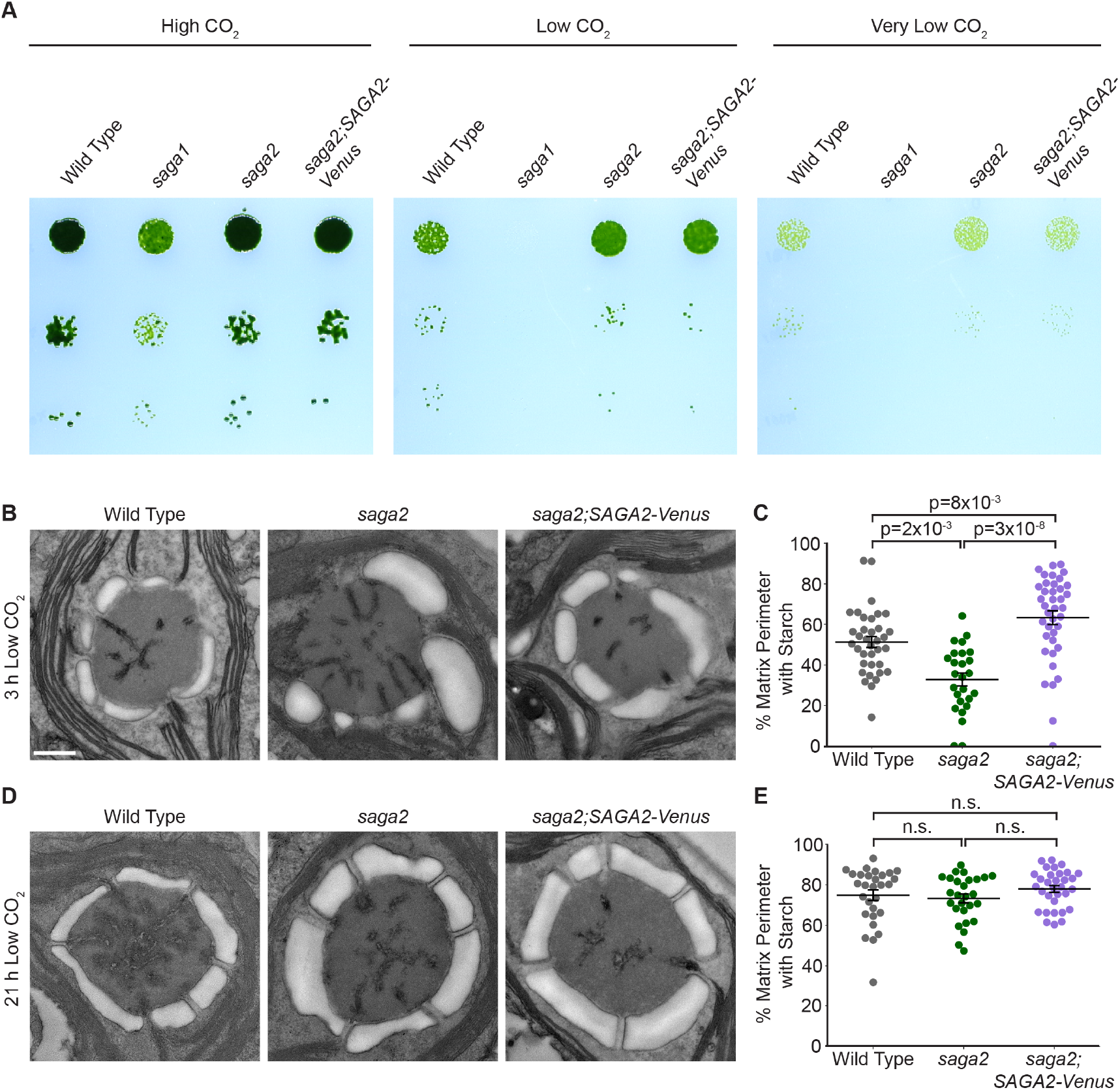
A *saga2* mutant grows normally under CO_2_-limiting conditions but has abnormal starch granules early after CO_2_-concentrating mechanism induction. (A) Agar growth phenotypes of wild type, *saga2*, and *saga2;*SAGA2-Venus. Serial 1:10 dilutions were spotted on TP minimal medium and grown at high (3% v/v), low (0.04% v/v), and very low (<0.004%) CO_2_ under 100 μmol photons⋅m^−2^⋅s^−1^ illumination. (B) Representative TEM images of wild-type, *saga2*, and *saga2;SAGA2-Venus* pyrenoids. Cells were grown at high (3% v/v) CO_2_ in minimal media, then moved to low (0.04% v/v) CO_2_ for 3 hours before harvesting. Scale bar = 500 nm. (C) Quantification of the proportion of the perimeter of the pyrenoid matrix in contact with starch in wild-type, *saga2*, and *saga2;SAGA2-Venus* pyrenoids at 3 hours low (0.04% v/v) CO_2_. (D) Representative TEM images of wild-type, *saga2*, and *saga2;SAGA2-Venus* pyrenoids. Cells were grown at high (3% v/v) CO_2_ in minimal media, then moved to low (0.04% v/v) CO_2_ for 21 hours before harvesting. (E) Quantification of the proportion of the perimeter of the pyrenoid matrix in contact with starch in wild-type, *saga2*, and *saga2;SAGA2-Venus* pyrenoids at 21 hours low (0.04% v/v) CO_2_. Statistical p-values were calculated using Kruskal-Wallis (p=7×10^−8^ in C, p=3×10^−1^ in E) followed by Dunn’s multiple comparisons test. Error bars represent the standard error of the mean.

### Absence of SAGA2 causes starch sheath defects early in pyrenoid formation

To determine whether the *saga2* mutant has starch sheath defects, we performed transmission electron microscopy (TEM) to visualize its starch sheath morphology. To induce starch sheath biogenesis, we transferred cells from high levels of CO_2_ to low levels of CO_2_ 3 hours before they were harvested and prepared for TEM. When we imaged these cells, we observed that starch was associated with the Rubisco matrix in both wild-type and *saga2* pyrenoids, but that the percentage of matrix perimeter covered with starch was lower in the *saga2* mutant than in wild type (33% in *saga2* vs 51% in wild type; p=2×10^−3^, Dunn’s multiple comparisons test; Fig. 2 *B* and *C* and Fig. S4*A*).

To confirm that this phenotype was caused specifically by the insertion in the *SAGA2* gene, we rescued the *saga2* mutant with *SAGA2-Venus* expressed under the native *SAGA2* promoter (Fig. 2*A*, Fig. S1*B*, and Fig. S2*B*). *SAGA2-Venus* rescued the *saga2* mutant phenotype; in the rescued strain, a greater percentage of matrix was covered with starch compared to the *saga2* mutant (63% in *saga2;SAGA2-Venus* vs 33% in *saga2*; p=3×10^−8^, Dunn’s multiple comparisons test; Fig. 2 *B* and *C* and Fig. S4*A*). Interestingly, the *saga2;SAGA2-Venus* strain also had more starch-matrix coverage than wild type (63% in *saga2;SAGA2-*Venus vs 51% in wild type; p=8×10^−3^, Dunn’s multiple comparisons test; Fig. 2 *B* and *C*, Fig. S4*A*, and Fig. S5), and the starch granules around the pyrenoids in *saga2;SAGA2-Venus* were thicker compared to those in wild type (Fig. 2*B* and Fig. S4*A*), which could potentially be due to overexpression of SAGA2-Venus in the *saga2;SAGA2-Venus* strain. Taken together, these results indicate that SAGA2 is necessary for normal matrix-starch sheath coverage during early stages of pyrenoid formation.

### The *saga2* mutant recovers a normal-looking starch sheath at later stages of pyrenoid maturation

The pyrenoid is a dynamic organelle that changes morphology over time after exposure to low CO_2_ levels (33–35). One such change is that the starch sheath in wild-type Chlamydomonas becomes thicker and expands to cover the full perimeter of the Rubisco matrix after the pyrenoid is fully induced (33, 35, 36). To determine whether these changes also occur in the *saga2* mutant, we prepared cells for TEM at a second time point 21 hours after we transferred cells from high CO_2_ to low CO_2_.

At the 21-hour time point, we indeed observed that the starch sheath in the *saga2* mutant was thicker, more tightly shaped around the Rubisco matrix, and covered a larger percentage of the matrix perimeter compared to the 3-hour time point (Fig. 2*B*-*E* and Fig. S4). For both the *saga2* mutant and *saga2;SAGA2-Venus*, the starch-matrix coverage of the pyrenoids showed no discernible difference compared to wild type at the 21-hour time point (p=3×10^−1^, Kruskal-Wallis ANOVA test; Fig. 2 *D* and *E* and Fig. S4*B*). Together, our findings suggest that while SAGA2 is necessary for a normal starch sheath early during pyrenoid formation, other factors can compensate for a lack of SAGA2 to assemble an apparently normal starch sheath during later stages of pyrenoid formation.

### The *saga2* mutant has a single pyrenoid and apparently normal tubules

Since SAGA1 was previously shown to be necessary for normal pyrenoid number and for pyrenoid tubule formation (18, 29), we sought to determine whether SAGA2, as its homolog, was also involved in these aspects of pyrenoid biogenesis. To determine whether the *saga2* mutant had a single pyrenoid, we transformed the *saga2* mutant with *Rubisco-mCherry* and imaged this strain along with a wild-type strain expressing *Rubisco-mCherry* (1, 29) both 3 hours and 21 hours after transferring cells to low CO_2_ using confocal microscopy. We found that both wild type and *saga2* had a single Rubisco punctus at the base of the chloroplast at both the 3-hour and the 21-hour time points (Fig. 3*A* and Fig. S6). We conclude that, unlike SAGA1, SAGA2 is not necessary for maintaining a single pyrenoid in Chlamydomonas.

**Figure 3.**
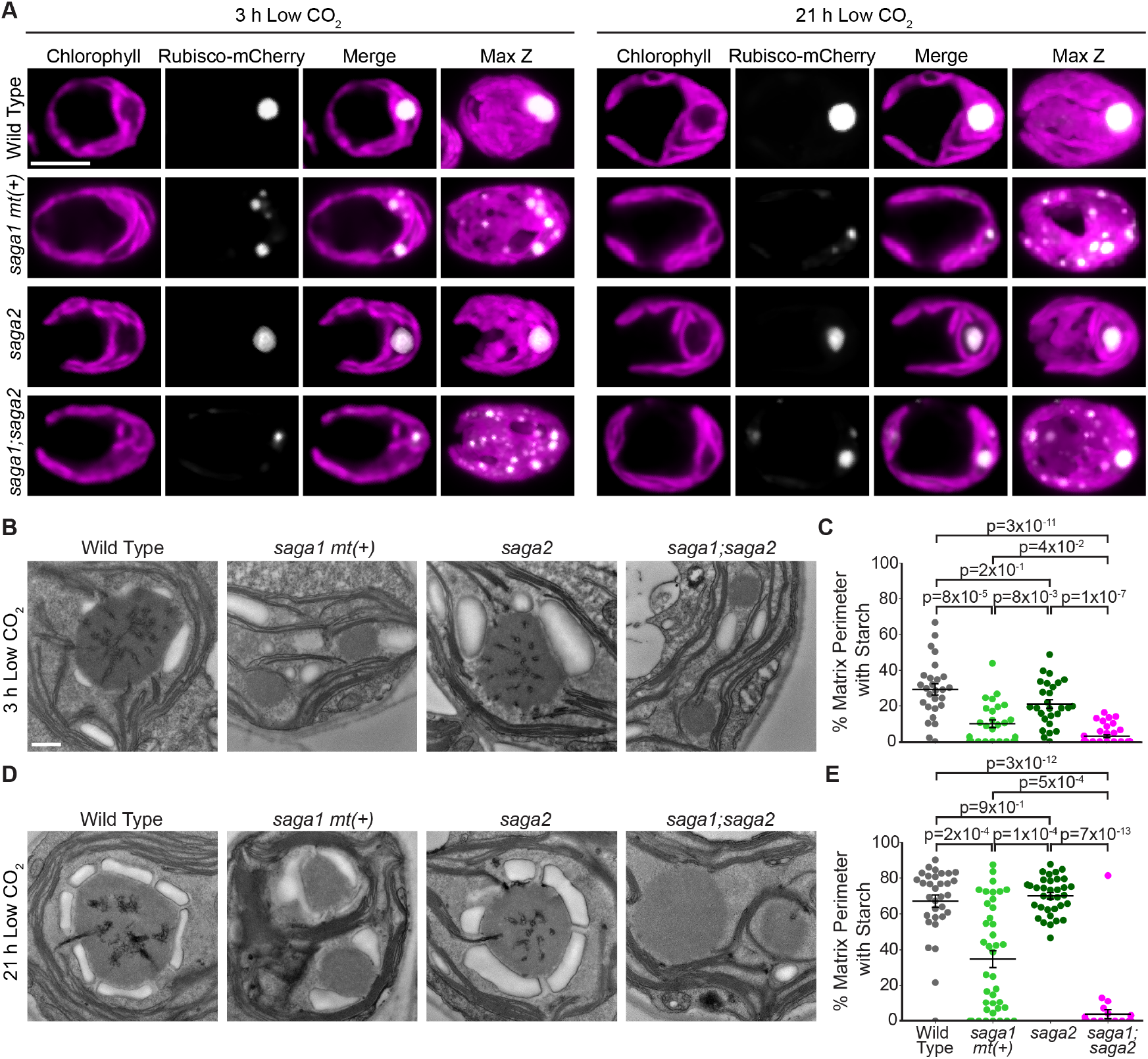
A *saga1;saga2* double mutant has multiple pyrenoid matrix condensates that lack starch sheaths. (A) Representative confocal images of RBCS1-mCherry in wild type, s*aga1 mt(+), saga2*, and *saga1;saga2* grown at high (3% v/v) CO_2_ in minimal media, then moved to low (0.04% v/v) CO_2_ for 3 hours (left) and 21 hours (right). Scale bar = 5 µm. (B) Representative TEM images of wild-type, *saga1 mt(+), saga2*, and *saga1;saga2* pyrenoids. Cells were grown at high (3% v/v) CO_2_ in minimal media, then moved to low (0.04% v/v) CO_2_ for 3 hours before harvesting. Scale bar = 500 nm. (C) Quantification of the proportion of the perimeter of the pyrenoid matrix in contact with starch in wild-type, *saga1 mt(+), saga2*, and *saga1;saga2* pyrenoids at 3 hours low (0.04% v/v) CO_2_. (D) Representative TEM images of wild-type, *saga1 mt(+), saga2*, and *saga1;saga2* pyrenoids. Cells were grown at high (3% v/v) CO_2_ in minimal media, then moved to low (0.04%) CO_2_ for 21 hours before harvesting. Scale bar = 500 nm. (E) Quantification of the proportion of the perimeter of the pyrenoid matrix in contact with starch in wild type, *saga1 mt(+), saga2*, and *saga1;saga2* pyrenoids at 21 hours low CO_2_. Statistical p-values were calculated with Kruskal-Wallis (p=1×10^−12^ in C, p=4×10^−16^ in E) followed by Dunn’s multiple comparisons test. Error bars represent the standard error of the mean.

To investigate whether SAGA2 is necessary for pyrenoid tubule formation, we analyzed the tubule networks of pyrenoids in our TEM images (Fig. 2 *B* and *D*, Fig. S4). At both the 3-hour and the 21-hour time points, all *saga2* mutant pyrenoids that we imaged had visible tubule networks that were similar to those we observed in wild type (Fig. 2 *B* and *D* and Fig. S4). These observations indicate that SAGA2 is not necessary for pyrenoid tubule formation.

### A *saga1;saga2* double mutant has multiple pyrenoid matrix condensates that lack thylakoid tubules

We next sought to investigate the epistatic relationship of SAGA1 and SAGA2 with respect to their impact on the starch sheath, pyrenoid condensate number, and tubule formation. To do this, we crossed a *saga1* mating type plus (*mt(+)*) strain (18) with the *saga2* mutant and isolated a *saga1;saga2* double mutant from the progeny (Fig. S7*A*-*F*).

To determine the number of pyrenoid matrix condensates in the *saga1;saga2* mutant, we transformed *Rubisco-mCherry* into *saga1 mt(+)* and *saga1;saga2* and imaged these strains along with *saga2;Rubisco-mCherry* both 3 hours and 21 hours after transferring cells to low CO_2_ using confocal microscopy. We found that both *saga1 mt(+)* and *saga1;saga2* had multiple Rubisco puncta distributed throughout the chloroplast, while the *saga2* mutant had a single Rubisco punctus at both the 3-hour and 21-hour time points (Fig. 3*A* and Fig. S6), indicating that the *saga1* mutation is epistatic to the *saga2* mutation with respect to pyrenoid condensate number.

To investigate whether *saga1;saga2* has normal pyrenoid tubules, we performed TEM on cells harvested at the 3-hour and 21-hour low CO_2_ time points. At both time points, we observed multiple small pyrenoid matrix condensates in *saga1 mt(+)* and *saga1;saga2*, visible as electron-dense regions with the same texture as the pyrenoid matrix (Fig. 3 *B* and *D*, Fig. S8, and Fig. S9). We found that, while *saga2* mutant pyrenoids had tubules, *saga1 mt(+)* and *saga1;saga2* both lacked visible thylakoid tubules in their multiple pyrenoid condensates (Fig. 3 *B* and *D*, Fig. S8, and Fig. S9). We also performed spot test assays to determine whether *saga1;saga2* had a functional pyrenoid and found that it had a severe growth defect similar to *saga1 mt(+)* at both low and very low CO_2_ levels (Fig. S3). Taken together, these results indicate that the *saga1* mutation is epistatic to *saga2* with respect to both pyrenoid matrix number and tubule formation, which likely causes the severe *saga1*-like growth defects observed in the *saga1;saga2* double mutant.

### Pyrenoid matrix condensates in the *saga1;saga2* double mutant lack starch sheaths

To investigate the relationship between SAGA1 and SAGA2 with respect to the starch sheath, we analyzed the pyrenoid starch sheaths in our TEM images of cells harvested at the 3-hour and 21-hour low CO_2_ time points. Remarkably, at both time points, the *saga1;saga2* mutant completely lacked starch sheaths around its visible pyrenoid condensates (Fig. 3*B-E*, Fig. S8, and Fig. S9), suggesting that SAGA1 and SAGA2 function redundantly to localize starch sheaths to the pyrenoid.

To validate that the lack of pyrenoid-associated starch sheaths in *saga1;saga2* was caused by the absence of SAGA1 and SAGA2, rather than a background mutation, we genetically rescued the *saga1;saga2* double mutant by transforming it with either *SAGA1-Venus* or *SAGA2-Venus* to generate *saga1;saga2;SAGA1-Venus* and *saga1;saga2;SAGA2-Venus*. We then visualized starch in the parental and rescued strains using fluorescein diacetate, a fluorescent dye that binds to starch (37), 3 hours and 21 hours after transferring cells to low CO_2_ (Fig. S10*A-E*). Canonical pyrenoid starch sheaths were visible in *saga1;saga2;SAGA1-Venus* at both time points, while multiple starch sheath rings were visible in *saga1;saga2;SAGA2-Venus* only at the 21-hour time point (Fig. S10*A-E*), which aligned with our TEM observations of starch sheaths in *saga2* and *saga1 mt(+)* (Fig. 2*B-E*, Fig. 3*B-E*, Fig. S4, and Fig. S8). Together, these findings establish that SAGA1 and SAGA2 can each compensate for the loss of the other to localize starch sheaths to the pyrenoid, but no other factor can compensate for the loss of both SAGA1 and SAGA2.

### The *saga1;saga2* double mutant produces less starch than wild type

Considering that starch is normally primarily found around the pyrenoid condensate when wild-type cells are grown under low CO_2_ (33), the absence of starch around the pyrenoid condensates in the *saga1;saga2* double mutant could indicate a complete lack of starch production in this strain. To determine whether the *saga1;saga2* mutant still produces starch, we used an iodine stain assay to detect starch in cell extracts from wild type, *saga1 mt(+), saga2, saga1;saga2*, and a starchless *sta6* mutant (38). This assay uses Lugol’s iodine solution, which turns blue when it binds to amylose and reddish-brown when it binds to amylopectin, the two main components of starch (39). We found that wild type, *saga1 mt(+), saga2*, and *saga1;saga2* all stained dark, while the *sta6* cell extract remained green (Fig. 4*A*). These results indicate that the *saga1;saga2* double mutant still produces starch, and that the absence of a starch sheath in *saga1;saga2* is not due to a complete lack of starch production.

**Figure 4.**
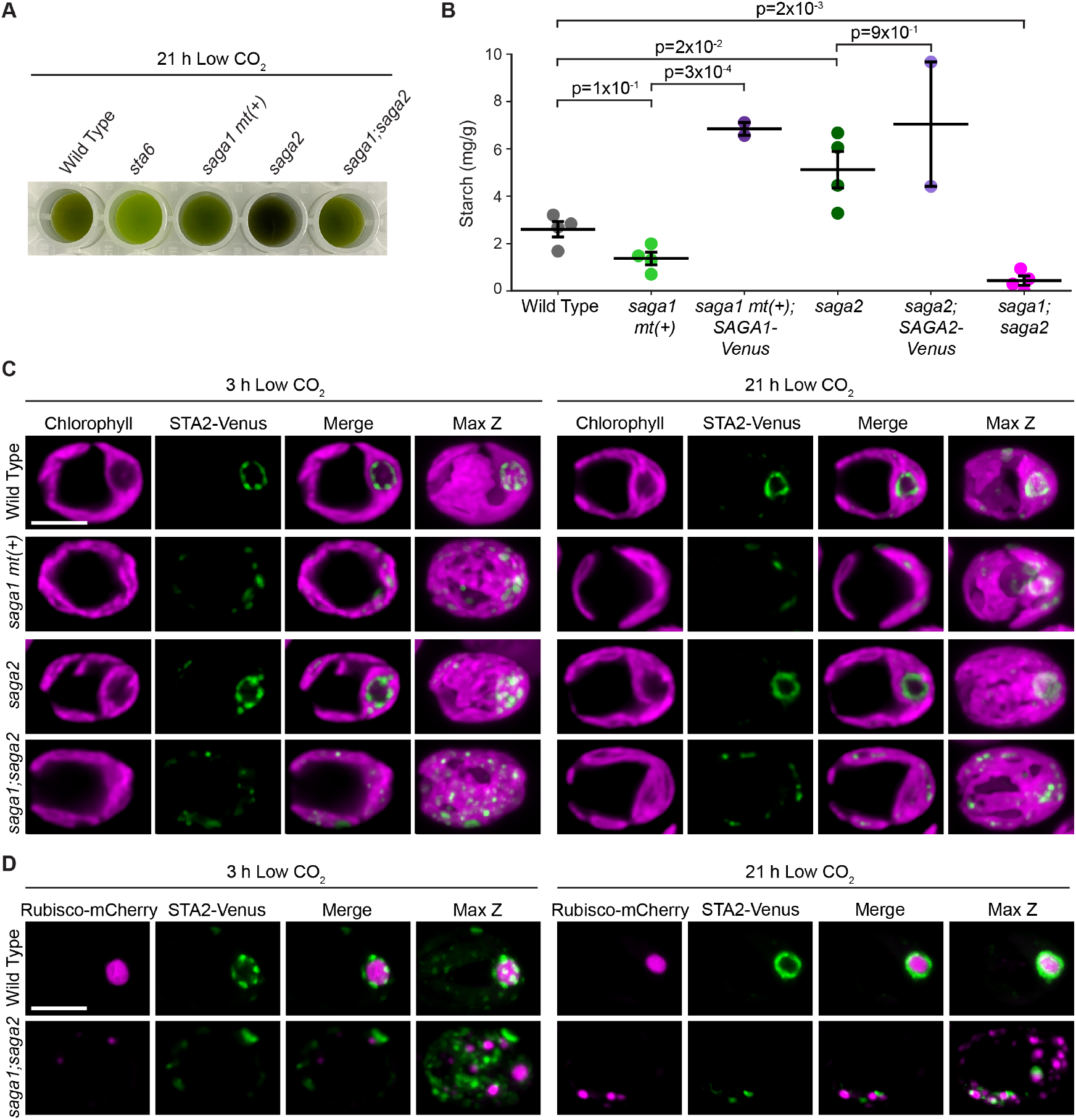
The *saga1;saga2* double mutant produces less starch than wild type, and its starch does not associate with the pyrenoid matrix. (A) Cell extracts from wild type, *sta6, saga1 mt(+), saga2*, and *saga1;saga2* stained with Lugol’s iodine solution. Cells were grown at high (3% v/v) CO_2_ in minimal media, then moved to low (0.04% v/v) CO_2_ for 21 hours before harvesting. (B) Quantification of the amount of starch produced in wild type, *saga1 mt(+), saga2, saga1;saga2, saga1 mt(+);SAGA1-Venus*, and *saga2;SAGA2-Venus*. Cells were grown at high (3% v/v) CO_2_ in minimal media, then moved to low (0.04% v/v) CO_2_ for 21 hours before harvesting. Statistical p-values were calculated using Kruskal-Wallis (p=4×10^−3^) followed by the Conover-Iman multiple comparisons test. Error bars represent the standard error of the mean. (C) Representative confocal images of STA2-Venus in wild type, s*aga1 mt(+), saga2*, and *saga1;saga2* grown at high CO_2_ in minimal media, then moved to low CO_2_ for 3 hours (left) and 21 hours (right) before imaging. Scale bar = 5 µm. (D) Representative confocal images of RBCS1-mCherry and STA2-Venus in wild type and *saga1;saga2* grown at high (3% v/v) CO_2_ in minimal media, then moved to low (0.04% v/v) CO_2_ for 3 hours (left) and 21 hours (right). Scale bar = 5 µm.

To determine whether there were differences in total starch content between the mutants, we purified starch from wild type, *saga1 mt(+), saga1 mt(+);SAGA1-Venus, saga2, saga2;SAGA2-Venus*, and *saga1;saga2*, then quantified how much starch was in each strain using a starch assay kit (Sigma). We found that *saga1 mt(+)* produced slightly less starch than wild type, although this change was not statistically significant (p=1×10^−1^, Conover-Iman multiple comparisons test; Fig. 4*B*). The *saga1 mt(+);SAGA1-Venus* rescue strain produced significantly more starch than *saga1 mt(+)* (p=3×10^−4^, Conover-Iman multiple comparisons test; Fig. 4*B*), suggesting that SAGA1 impacts starch production. The *saga2* mutant produced slightly more starch than wild type (p=2×10^−2^, Conover-Iman multiple comparisons test; Fig. 4*B*), but the *saga2;SAGA2-Venus* rescue strain produced a similar amount of starch to the *saga2* mutant (p=9×10^−1^, Conover-Iman multiple comparisons test; Fig. 4*B*). Strikingly, the *saga1;saga2* double mutant on average produced only 17% as much starch wild type (p=2×10^−3^, Conover-Iman multiple comparisons test; Fig. 4*B*). We conclude that the *saga1;saga2* double mutant produces less starch than wild type.

### Starch in the *saga1;saga2* double mutant is distributed throughout the stroma

Given that *saga1;saga2* had almost no matrix-associated starch granules but still produced some starch, we next sought to determine where that starch was localized. We used a fluorescently-tagged pyrenoid starch sheath marker, granule-bound starch synthase IA (40) (STA2, Cre17.g721500), to visualize the localization of starch 3 hours and 21 hours after transferring cells to low CO_2_ (Fig. 4*C* and Fig. S11). At both time points, in wild type and the *saga2* mutant, STA2-Venus localized in a well-defined ring around the pyrenoid (Fig. 4*C* and Fig. S11). In *saga1 mt(+)* at the 3-hour time point, STA2-Venus was distributed in puncta throughout the chloroplast, but at the 21-hour time point small rings were visible in the chloroplast in many cells, suggesting that starch sheaths had formed (Fig. 4*C* and Fig. S11). In *saga1;saga2* at both time points, STA2-Venus localized in puncta throughout the chloroplast (Fig. 4*C* and Fig. S11), with no visible ring-like patterns indicative of a pyrenoidal starch sheath. These results were validated by staining starch in *saga1;saga2* with fluorescein (Fig. S10*F*). Dual localization of *STA2-Venus* and *Rubisco-mCherry* further indicated that starch was predominantly not colocalized with the Rubisco condensates in the *saga1;saga2* mutant (Fig. 4*D* and Fig. S12). We conclude that in the *saga1;saga2* double mutant, starch is distributed throughout the stroma rather than to the pyrenoid matrix condensates.

### SAGA1 is expressed at similar levels in both wild type and *saga2*, and SAGA2 is present in *saga1*

Having established the impact of the *saga1* and *saga2* mutations on starch localization, we next sought to understand how SAGA1 and SAGA2 contribute to pyrenoid starch sheath localization in each other’s absence. For SAGA1 and SAGA2 to be able to compensate for each other’s loss, each protein would need to be present in the absence of the other. To investigate SAGA1 and SAGA2 protein levels in the mutants, we attempted to raise antibodies against SAGA1 and SAGA2 but failed to produce antibodies that were specific to each protein. Instead, we performed a western blot with an antibody that binds to both SAGA1 and SAGA2 (27) on cell extracts from wild type, *saga1 mt(+), saga1 mt(+);SAGA1-Venus, saga2, saga2;SAGA2-Venus*, and *saga1;saga2* (Fig. 5*A*). We found that SAGA1 levels were similar in wild type, *saga2*, and *saga2;SAGA2-Venus*. We cannot draw quantitative conclusions about the levels of SAGA2 in each strain because the SAGA2 band is faint and its intensity in different strain backgrounds varied between replicates. We conclude that SAGA1 protein levels are unchanged in the *saga2* mutant, and that SAGA2 is expressed in the *saga1* mutant, consistent with the idea that SAGA1 and SAGA2 can compensate for each other’s loss.

**Figure 5.**
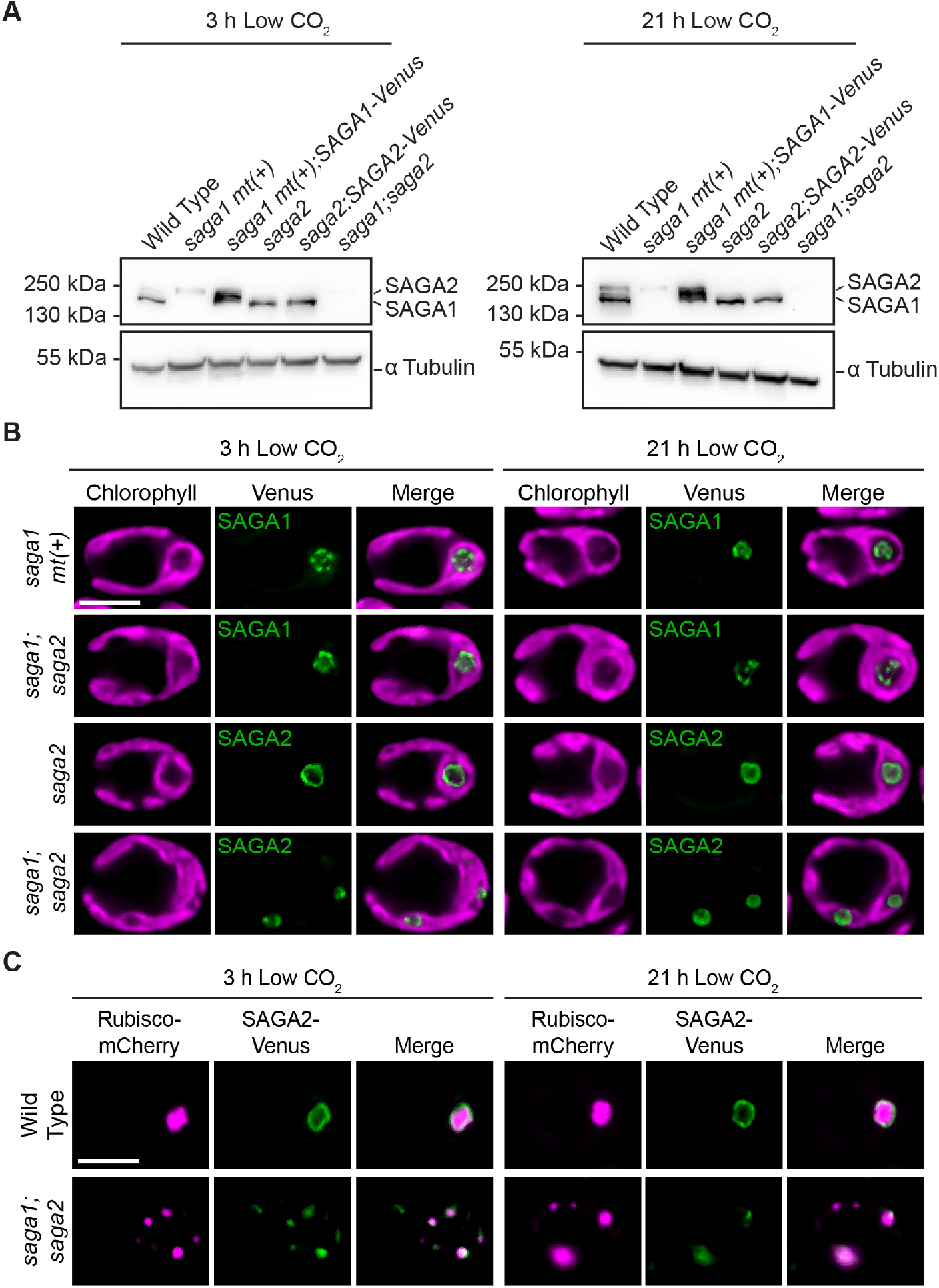
SAGA1 and SAGA2 are each expressed and localize to the periphery of the matrix in the absence of the other. (A) Anti-SAGA1 western blot of wild type, *saga1 mt(+), saga1 mt(+);SAGA1-Venus, saga2, saga2;SAGA2-Venus*, and *saga1;saga2* were grown at high (3% v/v) CO_2_ in minimal media, then moved to low (0.04% v/v) CO_2_ for 3 hours (top) and 21 hours (bottom) before harvesting. The antibody also appears to recognize SAGA2, as observed previously, but may not recognize SAGA2-Venus-3xFLAG, potentially due to the C-terminal tag, as has been observed for other proteins (18). (B) Representative confocal images of SAGA1-Venus in *saga1 mt(+)* and *saga1;saga2* and SAGA2-Venus in *saga2* and *saga1;saga2* grown at high (3% v/v) CO_2_ in minimal media, then moved to low (0.04% v/v) CO_2_ for 3 hours (left) and 21 hours (right). Scale bar = 5 µm. (C) Representative confocal images of Rubisco-mCherry and SAGA2-Venus in wild type and *saga1;saga2* grown at high (3% v/v) CO_2_ in minimal media, then moved to low (0.04% v/v) CO_2_ for 3 hours (left) and 21 hours (right). Scale bar = 5 µm.

### SAGA1 and SAGA2 each localize to the matrix periphery in the absence of the other

For SAGA1 and SAGA2 to be able to compensate for each other’s loss, we would expect each protein to localize to the matrix-starch interface in the other’s absence. As expected, SAGA1-Venus localized to its canonical location in both *saga1 mt(+)* and *saga1;saga2* (Fig. 5*B* and Fig. S13), and SAGA2-Venus localized in a single ring in the canonical pyrenoid location in the *saga2* mutant but localized to multiple puncta throughout the chloroplast in the *saga1;saga2* mutant (Fig. 5*B* and Fig. S13).

To determine whether SAGA2 was localized to multiple pyrenoid matrix condensates in the *saga1* mutant background, we imaged both SAGA2-Venus and Rubisco-mCherry in *saga1;saga2* (Fig. 5*C* and Fig. S14). Notably, SAGA2-Venus colocalized with some, but not all, Rubisco-mCherry puncta (Fig. 5*C* and Fig. S14), suggesting that SAGA2 only localized to a subset of the multiple pyrenoid matrix condensates in the *saga1* mutant background. Taken together, these results indicate that SAGA1 and SAGA2 can each localize to the matrix periphery in the absence of the other, and that SAGA2 associates with only a subset of the multiple matrix condensates of the *saga1* mutant.

### The SAGA1 and SAGA2 CBM20 domains bind to starch and the starch precursor maltoheptaose

To further understand how SAGA1 and SAGA2 can each localize starch to the pyrenoid, we sought to determine whether SAGA1 and SAGA2 can directly interact with starch and/or starch precursor molecules through their predicted CBM20 starch-binding domains (SBDs) (27, 29) (Fig. 1*B* and Fig. S15). We performed a starch-binding assay using purified CBM20 domains from either SAGA1 or SAGA2. To increase the sensitivity of the binding assay for SAGA1, we designed a construct with two copies of the SAGA1 CBM20 domain (10xHis-mVenus-2xSAGA1-SBD; Fig. S15). SAGA2 natively has two consecutive CBM20 domains, both of which were included in the SAGA2 construct for this experiment (10xHis-mVenus-SAGA2-SBD; Fig. S15).

We purified the starch-binding domains and assessed their binding to waxy maize starch granules in the presence or absence of two known CBM20 ligands, β-cyclodextrin and maltoheptaose (41, 42). We observed that both the 10xHis-mVenus-2xSAGA1-SBD and the 10xHis-mVenus-SAGA2-SBD proteins, but not a 10xHis-mVenus control protein, bound to starch (Fig. 6*A* and Fig. S16). Binding of 10xHis-mVenus-2xSAGA1-SBD and the 10xHis-mVenus-SAGA2-SBD to starch was decreased in the presence of either β-cyclodextrin or maltoheptaose, suggesting that each protein bound to all three molecules. β-cyclodextrin is not produced by Chlamydomonas cells, but is a soluble cyclic seven-residue glucose polymer that mimics the helical conformation of starch as well as that of long maltooligosaccharides, which are substrates for starch initiation (42–44); maltoheptaose is a linear seven-residue maltooligosaccharide that can act as a primer for the initiation of starch synthesis (43). The binding of 2xSAGA1-SBD and SAGA2-SBD to both β-cyclodextrin and maltoheptaose suggests that SAGA1 and SAGA2 can bind to both linear and helical starch precursor molecules. For both 2xSAGA1-SBD and SAGA2-SBD, β-cyclodextrin decreased binding more than maltoheptaose (Fig. 6*A* and Fig. S16), suggesting that SAGA1 and SAGA2 may bind more strongly to helical starch precursors. We conclude that both the SAGA1 CBM20 domain and the SAGA2 CBM20 domain can bind to not only starch but also to starch precursor molecules, suggesting that SAGA1 and SAGA2 could promote starch initiation at the surface of the pyrenoid.

**Figure 6.**
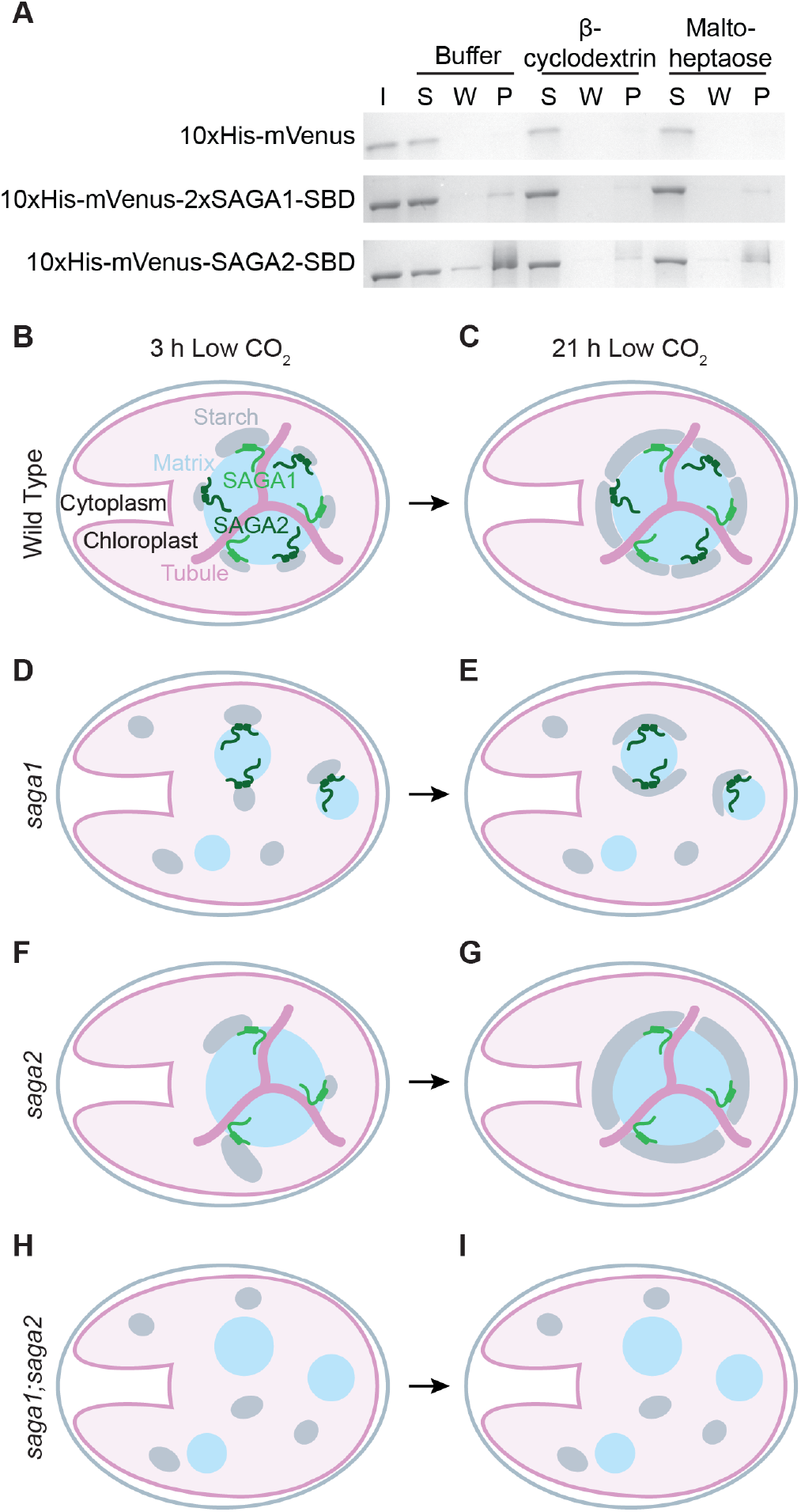
The SAGA1 and SAGA2 CBM20 domains bind starch and the starch precursor maltoheptaose, suggesting a role in starch sheath initiation. (A) Starch binding assay of the predicted SAGA1 and SAGA2 CBM20 starch binding domains (SBDs) with starch in the presence or absence of the starch mimic β-cyclodextrin and the starch precursor maltoheptaose. The purified proteins were incubated with starch granules in either buffer or buffer supplemented with β-cyclodextrin or maltoheptaose. The starch granules were then pelleted, washed three times, and the protein that remained bound to starch was solubilized and denatured before loading onto an SDS-PAGE gel. I – input protein in buffer. S – supernatant after initial incubation with starch. W – the final wash. P – protein that remained bound to the starch pellet. (B-I) A proposed model for how SAGA1 and SAGA2 localize starch around the pyrenoid. (B), (C) In wild type, SAGA1 and SAGA2 each initiate starch granules at the pyrenoid periphery. (D), (E) In the *saga1* mutant, only SAGA2 initiates granules at the surface of the pyrenoid condensates. (F), (G) In the *saga2* mutant, only SAGA1 initiates granules at the pyrenoid periphery. (H), (I) In the *saga1;saga2* double mutant, starch granules are not initiated in proximity to the matrix, and the starch is not shaped into a sheath. Instead, starch granules initiate in the stroma away from the matrix condensates. In all but the *saga1;saga2* double mutant, initiated granules grow to cover the condensates they are associated with.

## Discussion

### Our data suggest that SAGA1 and SAGA2 are starch granule initiation factors

SAGA1 and SAGA2 were previously suggested to be matrix-starch tethers (27), in which case other starch biogenesis factors would initiate starch granules at the surface of the matrix, and SAGA1 and SAGA2 would tether this starch to the matrix. However, our data raise the possibility that SAGA1 and SAGA2 do not merely tether starch to the matrix, but rather directly promote starch granule initiation at the surface of the pyrenoid.

The first piece of evidence that SAGA1 and SAGA2 are initiators rather than simply tethers is that the *saga1;saga2* double mutant lacks matrix-associated starch granules. If factors other than SAGA1 and SAGA2 were responsible for initiating starch granules at the matrix periphery, we would naïvely expect that the initiated granules would remain in the proximity of the matrix even in the absence of tethers – but instead, in the *saga1;saga2* double mutant, we observe starch distributed throughout the stroma (Fig. 3 *B* and *D*, Fig. 4 *C* and *D*, Fig. S8, Fig. S9, Fig. S10*F*, Fig. S11, and Fig. S12). This observation suggests that starch in the *saga1;saga2* double mutant was initiated away from the matrix and that the double mutant lacks the ability to initiate starch at the matrix periphery, implicating SAGA1 and SAGA2 in starch granule initiation.

The second piece of evidence that SAGA1 and SAGA2 are starch initiators is that the *saga1;saga2* double mutant produces less starch than wild type (Fig. 4*B*). If SAGA1 and SAGA2 were merely acting as tethers, we would not necessarily expect to see an impact on the total amount of starch produced by the cell. By contrast, if SAGA1 and SAGA2 are starch granule initiators, the decrease in total starch in the *saga1;saga2* double mutant can be explained by fewer starch initiation events.

Another piece of data consistent with the idea that SAGA1 and SAGA2 could be starch initiators is that SAGA1 remains localized in a punctate pattern around the pyrenoid in the absence of SAGA2 (Fig. 5*B* and Fig. S13) even though the starch sheath fully surrounds *saga2* pyrenoid condensates (Fig. 2 *D* and *E*, Fig. 3 *D* and *E*, Fig. S4, and Fig. S8). This observation indicates that SAGA1 only needs to act at puncta to achieve full starch sheath coverage of the matrix, which is consistent with an initiation function for SAGA1. Furthermore, the observation that starch sheaths are in contact with the matrix along the full surface of the pyrenoid in the *saga2* mutant (Fig. 2 *D* and *E*, Fig. S4, and Fig. S8) suggests that factors other than SAGA2 promote the close apposition of starch with the matrix at sites distant from SAGA1.

A final piece of evidence that supports the idea that SAGA1 and SAGA2 are starch initiators is that SAGA1 and SAGA2 share domain, sequence, and functional similarities with established Arabidopsis starch initiation factors. While the mechanisms for starch initiation in Chlamydomonas remain poorly understood, more is known about this process in Arabidopsis, where starch granules are initiated in stromal pockets between thylakoid membranes in the chloroplast (45). The specific sites for starch initiation are thought to be determined by the localization of the thylakoid-bound coiled-coil protein MAR BINDING FILAMENT-LIKE PROTEIN 1 (MFP1, AT3G16000), which binds to the non-enzymatic starch initiator PROTEIN TARGETING TO STARCH 2 (PTST2, AT1G27070) (45, 46). PTST2 helps initiate starch by binding both to maltooligosaccharides (MOS) and to SOLUBLE STARCH SYNTHASE 4 (SS4), bringing SS4 into contact with its MOS substrates to enable starch granule initiation (43). While there are no detectable Chlamydomonas homologs of PTST2 (47), SAGA1 and SAGA2 both share predicted structural similarity to PTST2, which is also a coiled-coil CBM-containing protein (43, 45). Additionally, SAGA1 and SAGA2 share weak sequence similarity to MFP1 homologs in several cereal species (Fig. S17 and Table S2). Furthermore, we show that, like PTST2, SAGA1 and SAGA2 can bind to MOS (Fig. 6*A* and Fig. S16) (43).

The similarities of SAGA1 and SAGA2 with MFP1 and PTST2, along with our evidence that SAGA1 and SAGA2 are likely involved in starch initiation, lead us to speculate that SAGA1 and SAGA2 could perform the combined function of MFP1 and PTST2. Rather than establishing the site of starch initiation in stromal pockets, as MFP1 is proposed to do, SAGA1 and SAGA2 could establish the sites of starch initiation at the surface of the pyrenoid by binding to Rubisco via their Rubisco-binding motifs (Fig. 1 *A* and *B*) (27) and by binding MOS via their starch-binding domains (Fig. 1*B*). While we did not observe interactions between SAGA1 and SAGA2 and any starch synthase by immunoprecipitation-mass spectrometry (Table S3), such interactions may occur but be too transient to detect with our assays. Alternatively, if SAGA1 and SAGA2 do not directly interact with a starch synthase, they could promote starch granule initiation near the pyrenoid by increasing the relative concentration of MOS around the matrix. This idea is supported by recent work that found that an increase of MOS in the chloroplast promoted starch granule initiation in Arabidopsis (48).

### We propose a model for how SAGA1 and SAGA2 localize starch to the pyrenoid in Chlamydomonas

We propose a model for starch sheath initiation at the periphery of the pyrenoid that is consistent with our data and with the hypothesis that SAGA1 and SAGA2 initiate starch granules at the surface of the Rubisco matrix (Fig. 6 *B-I*). In wild-type Chlamydomonas, SAGA1 initiates starch synthesis by binding to MOS at the surface of the matrix near pyrenoid tubule entry sites. SAGA2, on the other hand, initiates starch synthesis by binding to MOS along the surface of the matrix between tubule entry sites (Fig. 6*B*). From these early initiation sites, starch granules then grow around the surface of the matrix to fully encapsulate the pyrenoid by 21 hours at low CO_2_ (Fig. 6*C*).

In the *saga1* mutant, SAGA2 initiates starch along the surface of the multiple pyrenoid matrix condensates (Fig. 6*D*). Because SAGA2 is dispersed across multiple pyrenoid condensates, each pyrenoid matrix has only a small number of starch initiation points, leading to a delay in pyrenoid starch initiation and ultimately less matrix-starch coverage at the 3-hour time point. In the *saga2* mutant, SAGA1 initiates starch at the surface of the Rubisco matrix near tubule entry sites (Fig. 6*F*). The lack of SAGA2 initiation points along the surface of the matrix leads to less matrix-starch coverage at the 3-hour time point. By later time points in each single mutant, the starch granules that each SAGA protein has initiated on its own have had enough time to elongate along the matrix surface to encapsulate the pyrenoid, resulting in normal starch sheath coverage (Fig. 6 *E* and *G*). In the *saga1* mutant, some matrix condensates have no observed starch sheath even after 21 hours at low CO_2_ (Fig. 3*D-E* and Fig. S8). This can be explained by our observation that SAGA2 only localizes to a subset of the pyrenoid matrix condensates in the *saga1* mutant (Fig. 5*C* and Fig. S14). Based on our data and model, we speculate that the condensates that lack SAGA2 are the ones that do not form starch sheaths.

In the *saga1;saga2* double mutant, the lack of both SAGA1 and SAGA2 results in no starch being initiated at the matrix surface, resulting in no starch sheath forming around the multiple pyrenoid matrix condensates (Fig. 6 *H* and *I*). This model is consistent with our results as well as with previous results that showed that SAGA1 and SAGA2 promote the recruitment of starch to engineered Rubisco condensates in Arabidopsis (31).

Interestingly, our work suggests that the canonical Chlamydomonas starch sheath shape is dependent on starch granules being in contact with the Rubisco matrix; in the absence of such contact, granules do not appear to form in the shape of a starch sheath (Fig. 3 *B* *and* *D*, Fig. 4 *C* and *D*, Fig. S8, Fig. S9, Fig. S10*F*, Fig. S11, and Fig. S12). Instead, starch in *saga1;saga2* appears similar in shape to typical stromal starch.

### The starch localization principles in Chlamydomonas may apply more broadly

While progress has been made in understanding the factors involved in pyrenoid assembly in Chlamydomonas, little is known about the assembly of pyrenoids in other algal species. Pyrenoids are thought to have evolved independently via convergent evolution (3, 49–51), and as such, pyrenoid-related functions in different algal lineages are not necessarily performed by homologous proteins. Known Chlamydomonas pyrenoid proteins, including SAGA1 and SAGA2, as well as the Chlamydomonas Rubisco-binding motif, are conserved among closely related members of the green algal order Volvocales, but are not found in other algal species (27). There is, however, growing evidence that algal species outside of the Volvocales evolved their own unique Rubisco-binding motifs and likely assemble their Rubisco matrices using linker proteins that perform a similar function to EPYC1 in Chlamydomonas (52, 53). While the exact protein sequences may have no homology, we speculate that other algal species assemble pyrenoid starch sheaths using functional homologs of the Chlamydomonas SAGA proteins. Such proteins would likely be structurally similar to SAGA1 and SAGA2, with carbohydrate-binding domains and copies of whichever Rubisco-binding motif evolved in that species. Thus, beyond advancing the understanding of pyrenoid starch sheath assembly in Chlamydomonas, this work contributes to the foundation for investigating starch sheath biogenesis in other algal species.

## Materials and Methods

### Strains and Culture Conditions

Strains used in this study can be found in Table S4. All strains were maintained at room temperature (RT) (∼22°C) under very low light (<10 μmol photons⋅m^−2^⋅s^−1^) on 1.5% agar plates containing Tris-acetate-phosphate (TAP) medium with revised trace elements (54) supplemented with carbendazim. For liquid cultures, strains were grown in an orbital shaking incubator (Infors) using standard settings of 23°C and 120 rpm under continuous cool white LED light at ∼175 μmol photons⋅m^−2^⋅s^−1^. Unless otherwise noted, cultures used in experiments were grown in Tris-Phosphate (TP, same composition as TAP above but without acetate) liquid media in air enriched to 3% (v/v) CO_2_ for 2-3 days. Cultures were then diluted and moved to a different incubator with the same parameters except at low (0.04% v/v) levels of CO_2_ for either 3 hours or 21 hours. Cultures were harvested for experiments once they grew to approximately 2 x 10^6^ cells⋅mL^−1^ as measured by a Countess II Automated Cell Counter (Life Technologies).

All strains generated in this work were deposited to the Chlamydomonas Resource Center (https://chlamycollection.org).

### Crossing of Chlamydomonas Strains

The *saga1;saga2* double mutant used in this study was produced through crossing the opposite mating-type strains *saga1 mt(+)* and *saga2 mt(-)* (Table S4). A modified version of a previously published protocol (55) was performed as described in Hennacy et al., 2024, with two exceptions: 1) equal volumes of *saga1 mt(+)* and *saga2 mt(-)* were left at ∼175 µmol photons m^−2^⋅s^−1^ light without agitation for 1, 2, 3, and 4 hours, with the positive *saga1;saga2* double mutant coming from the 3 hour time point; 2) because *saga1 mt(+)* and *saga2 mt(-)* have the same antibiotic resistance, antibiotic selection could not be used to select for the correct genotypes. Instead, colonies were selected at random and screened through PCR to determine the genotypes (Table S1).

### Cloning

Plasmids used in this study are listed in Table S5. The generation of these plasmids is described in the SI Materials and Methods. Constructs were validated using nanopore sequencing (Plasmidsaurus).

### Transformation of Chlamydomonas

The strains in this study produced through Chlamydomonas transformation are noted in Table S4. Leading up to the transformation protocol cells were grown as follows. To generate *saga2;RBCS1-mCherry* and *saga1;saga2;RBCS1-mCherry*, cells were grown in liquid TAP in low (0.04% v/v) CO_2_. To generate *saga2;SAGA2-Venus*, cells were grown in liquid TAP in low (0.04% v/v) CO_2_ for 1 day, then moved to high (3% v/v) CO_2_. For all other transformations, cells were grown in liquid TAP in 3% (v/v) CO_2_. All strains for transformation were grown with the standard settings described above.

All transformations were performed as described in Hennacy et al., 2024 with some exceptions as described in the SI Materials and Methods. Positive transformants were selected for on TAP 1.5% agar plates supplemented with carbendazim and the appropriate antibiotic (Table S4) and kept in low light until colonies were a sufficient size for picking. Plates supplemented with zeocin were left at dark ambient light for ∼6 days to allow time for the light-sensitive antibiotic to begin to kill negative transformants before exposing the cells to ∼100 μmol photons⋅m^−2^⋅s^−1^ light. Once colonies were large enough, they were selected at random either by hand or using a PIXL precision microbial colony picker (Singer Instruments). All positive transformants were confirmed using a Nikon A1R point scanning confocal microscope to detect transgene expression via fluorescence.

### Spot Tests

Cells were grown to ∼2 x 10^6^ cells⋅mL^−1^ in liquid TAP medium at low (0.04%) levels of CO_2_ using the standard settings described above. They were then harvested at 1,000 x g for 5 minutes and cell pellets were washed 1x with 5 mL liquid TP media. Cell pellets were then resuspended in liquid TP to a concentration of ∼6 x 10^5^ cells⋅mL^−1^. Each sample was serially diluted at 1:10 and 1:100 dilutions, and 10 μL of the original sample and each dilution were then spotted onto TP 1.5% agar plates supplemented with carbendazim. Spots were allowed to dry and then plates were placed under ∼100 μmol photons⋅m^−2^⋅s^−1^ of light at high (3% v/v), low (0.04% v/v), and very low (<0.004% v/v) CO_2_ levels for 7 days before imaging on a PhenoBooth imager (Singer Instruments).

### Fluorescein Staining

To visualize starch in different Chlamydomonas strains, 1 mL of exponential phase cultures were harvested by centrifugation at 1,000 x g for 5 minutes using a swinging bucket rotor. The supernatants were discarded and cell pellets were resuspended in 999 µL fresh TP medium before adding 1 µL 50 mM fluorescein diacetate (FDA). Cells were pulse-vortexed and incubated for 30 minutes at room temperature in a thermomixer set to 1,000 rpm. After 30 minutes cells were then prepared for confocal imaging. The 50 mM FDA solution was prepared as follows. 2.1 mg of FDA powder (Chem Impex 01474) was dissolved by vortexing briefly in 10 µL of 2% Pluronic F-127 (Sigma P2443) in DMSO until the solution appeared cloudy. Then, 90 µL of DMSO was added and the entire solution was vortexed in pulses until the powder dissolved.

### Confocal Microscopy

For most confocal experiments, live cells were imaged with a Nikon A1R point scanning confocal microscope using a 100x magnification oil objective with pinhole set to 1.2 Airy Units (AU), and images were denoised using the Nikon NIS-Elements Denoise.ai function and then display values were adjusted for appropriate contrast with Fiji software (56). To visualize fluorescence throughout the chloroplast, 0.3 μm Z-slices were collected from the top to the bottom of each cell and Max-Z projections were created using Fiji. For all mCherry-expressing strains, the mCherry channel was collected with an excitation/emission of 561 nm/570-620 nm and chlorophyll autofluorescence was collected at 561 nm/663-738 nm. For all Venus-expressing strains and fluorescein-stained cells, Venus and fluorescein fluorescence were collected with 514 nm excitation and 522-555 nm emission and chlorophyll autofluorescence was collected with 514 nm excitation and 601-676 nm emission.

For *saga1;SAGA1-Venus* and *saga2;SAGA2-Venus* images, live cells were imaged with a Nikon Ti2 Inverted Microscope with a Yokogawa W1 and SoRa Module using a 60x TIRF objective with 4x SoRa magnification. Venus fluorescence was collected with 514 nm excitation and a 545/30 nm emission filter and chlorophyll autofluorescence was collected with 640 nm excitation and a 700 nm long pass emission filter. An alignment offset between the 514 and 640 channels was corrected as described in the SI Materials and Methods. All SAGA1-Venus and SAGA2-Venus images were denoised using the NIS-Elements Denoise.ai function and underwent 2D Automatic deconvolution. The images were then adjusted to display appropriate contrast using Fiji software (56).

### Analysis of Colocalization of SAGA1-Venus and SAGA2-Venus with Chlorophyll Autofluorescence

To determine SAGA1-Venus and SAGA2-Venus fluorescence intensity relative to tubule chlorophyll autofluorescence intensity levels, a freehand circle was drawn through the center of the SAGA-Venus signal around the periphery of the pyrenoid in merged images using Fiji software (Fig. S1) (56). The Plot Profile command was then used to measure fluorescence intensity levels for each individual channel. These values were transferred into Excel and used to create radial line graphs for data visualization and to calculate correlation coefficients.

### Transmission Electron Microscopy

For TEM, cells were fixed with glutaraldehyde, stained with OsO_4_, and embedded in Quetol epoxy resin as described in Hennacy et al., 2024, with the exception that for the serial dehydration step, samples were incubated for 5 minutes in 30%, 50%, 70%, and 95% ethanol, followed by 10 minutes in 100% ethanol and finally twice for 10 minutes in 100% acetonitrile, pelleting at 3,000 x g between each incubation. Once cells were embedded in resin, the resin blocks were sectioned into ∼70 nm slices using ultramicrotomy with DiaTome diamond knives on a Leica UCT Ultramicrotome. Resin slices were then mounted on carbon film-coated 200 mesh copper TEM grids (Electron Microscopy Sciences) and the cells were imaged using a Talos L120C G2 TEM microscope.

### Analysis of Starch-Matrix Coverage in TEM Images

To determine the percent of the pyrenoid matrix in contact with starch in TEM images, all images were first anonymized to minimize bias. The anonymized images were then analyzed using Fiji (56) as follows. First, the perimeter of the pyrenoid matrix was traced with a freehand circle, and the length of the circumference was measured. Then, for each starch granule in contact with the matrix, the length of the portion of the matrix in contact with that starch granule was measured. After all images per dataset were traced and measured, the images were de-anonymized to determine which strain each set of measurements corresponded to. For each individual pyrenoid, the lengths of the portions of matrix in contact with starch were summed and divided by the length of the perimeter of the matrix and the number of pyrenoid starch granules was counted using Python. A Kruskal-Wallis ANOVA test followed by Dunn’s multiple comparisons test was used to test for significance in starch coverage between strains and grouped scatter plots were created for data visualization using Python (seaborn.swarmplot).

### Protein Immunoprecipitation and Mass Spectrometry

For immunoprecipitation and mass spectrometry of SAGA1-Venus-3xFLAG, SAGA2-Venus-3xFLAG, and Venus-3xFLAG, cells were grown in 1 L bottles in TP medium with air bubbling and constant stirring at 210 rpm under ∼150 μmol photons⋅m^−2^⋅s^−^1 light before collecting by centrifugation at 3,000 x g for 4 minutes at 4°C. The protocol for washing, storing, and lysing pellets and the following immunoprecipitation and mass spectrometry was performed as described previously (18, 57). The Venus-3xFLAG data in Table S3 was previously published as a control in Hennacy et al., 2024 (18).

### Western Blots

Cells for western blots were grown in liquid TP at air CO_2_ under the standard conditions described above. Cell suspensions were pelleted, and pellets were resuspended in 300 μL of lysis buffer (5 mM Hepes-KOH pH 7.5, 100 mM dithiothreitol, 100 mM Na_2_CO_3_, 2% SDS, 12% sucrose, and cOmplete protease inhibitor cocktail) before transferring to a 1.5 mL microcentrifuge tube. Samples were heat-denatured, and lysates were clarified before aliquoting and storing at −80°C until analysis with SDS-polyacrylamide gel electrophoresis. After running samples on a tris/glycine gradient gel (10%, Bio-Rad Mini-Protean TGX Precast Gel), proteins were transferred to a 0.45 μm polyvinylidene difluoride membrane (Immobilin-P, MilliporeSigma) using a semi-dry transfer system (Bio-Rad) and transfer buffer (20% v/v ethanol, 25 mM Tris, 192 mM glycine, and 0.05% SDS).

For immunoblot analysis, membranes were blocked in TBST (tris-buffered saline from Bio-Rad + 0.1% Tween-20 from Sigma) containing 5% non-fat dry milk (LabScientific). Incubations with the primary antibody (anti-SAGA1 from Meyer et al., 2020, 1:700 dilution) were performed in TBST containing 2.5% milk. Membranes were washed in TBST 3x before incubation with the secondary antibody (Goat anti-rabbit IgG H+L HRP, Invitrogen 31466, 1:3,000 dilution). Membranes were washed again 3x in TBST. Immunoreactive proteins were visualized using enhanced chemiluminescence (WesternBright ECL, Advansta) followed by detection using an iBright 1500 Imaging System (Invitrogen).

### Lugol’s Iodine Stain Assays

Iodine stain experiments were performed following a modified version of a previously published protocol (58). Cultures were grown as above and were transferred to low (0.04% v/v) CO_2_ 21 hours before harvesting once cells grew to ∼2 x 10^6^ cells⋅mL^−1^. Cells were centrifuged at 1,000 x g for 5 minutes and cell pellets were resuspended in 5 mL liquid TP. Cells were then sonicated on ice for 2 minutes (cycles of 6 second pulse at 60% amplification, 3 second rest) using a probe sonicator (QSonica). Lysates were then centrifuged at 2,000 x g for 20 minutes and pellets were resuspended with 5 mL Tris-HCl, pH 8.0. Samples were then centrifuged at 2,000 x g for 5 minutes and the pellets were resuspended in 500 μL Milli-Q water. 500 μL of each resuspension was then transferred to a 1.5 mL microcentrifuge tube and 50 μL of Lugol’s iodine solution (Sigma) was added. 100 μL of each sample was then transferred to a 96-well plate for imaging.

### Starch Purification and Quantification

To extract and purify starch from Chlamydomonas, a previously published protocol was adapted (58). Briefly, cells were grown in 1 L cultures with air bubbling and constant stirring of 230 rpm under ∼150 μmol photons⋅m^−2^⋅s^−1^. Cultures were harvested, pellets were weighed and resuspended 1:10 w/v in TP media, and equal volumes of each were transferred to a 5 mL Eppendorf tube. Cells were then lysed by sonication at 4ºC, the lysate was clarified, and the pellets were rinsed with 5 mL 10 mM Tris-HCl, pH 8.0. Pellets were rinsed and then loaded onto 5 mL 100% Percoll (Sigma P7828) and centrifuged at 3,428 x g for 20 minutes. Starch pellets were rinsed 2x with dI water, then stored at 4ºC overnight. Pellets were then boiled in dI water for 3 minutes and autoclaved for 1 hr at 121ºC. The solubilized starch was then quantified following the protocol for the Sigma Starch Assay Kit (SA-20).

### CBM20 Domain Protein Purification

The plasmids containing the 10xHis-mVenus-CBM20 sequences coding for the SAGA1 and SAGA2 CBM20 domains were expressed in BL21 (DE3) cells (NEB) using an autoinduction system as described in the SI Materials and Methods. Following growth and induction, cells were harvested and lysed using sonication, and the resulting lysate was clarified. The supernatant was then supplemented with 50 mM Imidazole (Thermo Scientific) and filtered using a 0.22 µm vacuum filter (Millipore) before affinity purification using a HisTrap HP column (Cytiva). The purified proteins were then buffer-exchanged into starch binding buffer to be used in the starch binding assay.

### Starch Binding Assay

To test whether the SAGA CBM20 domains bind starch, waxy maize starch (Sigma) was washed in starch binding buffer (50 mM HEPES pH 7.8, 25 mM KCl, 2 mM MgCl_2_, 5 mM TCEP, 5% [v/v] glycerol), then pelleted and blocked with 5% (w/v) BSA (AG Scientific) for 1 hour with gentle rotation using a tube rotator (VWR). The starch was pelleted again, then washed 3x with starch binding buffer. Purified CBM20 domains or the 10xHis-mVenus control in starch binding buffer or buffer supplemented with β-cyclodextrin (Sigma) or maltoheptaose (Sigma) were added to the starch and incubated for 1 hour with gentle rotation. Pellets were then washed 3x, then incubated in 1x Laemmli buffer (Bio-Rad) at 37°C with agitation. Samples were taken from the initial input protein, the supernatant prior to the first wash, the final wash, and from the Laemmli buffer elution, then loaded onto a Mini-Protean TGX Precast polyacrylamide gel (Bio-Rad) for visualization. To detect the mVenus fluorescence emitted from proteins bound to the starch pellets, the starch pellets before the first wash and after the final wash were imaged on an Invitrogen iBright 1500 using the fluorescent blot setting.

### Protein Alignments

SAGA1, SAGA2, and MFP1 amino acid sequences were aligned using BLASTP (blast.ncbi.nlm.nih.gov). For any two proteins that returned more than one possible alignment, the E-value, bit score, and percent identity from the best possible alignment were reported.

The SAGA1 and SAGA2 Interpro-predicted CBM20 domain sequences (59, 60) were aligned with the CBM20 domain sequence from *Aspergillus niger* glucoamylase G1 (PDB ID: 1ACZ) using Clustal Omega (61–63). AlphaFold 3 (64) was used to generate high confidence predicted protein structures for each SAGA CBM20 domain. These predicted structures were aligned to the NMR solution structure of the *A. niger* glucoamylase G1 CBM20 domain in complex with β-cyclodextrin at the starch-binding site (61, 65) and the NMR solution structure of the free *A. niger* glucoamylase G1 CBM20 domain without β-cyclodextrin (1KUL) (66, 67) using the RCSB (RCSB.org) PDB pairwise structure alignment software (68, 69). All molecular graphics were created using ChimeraX software (70).

## Acknowledgments

We thank Moritz T. Meyer for helpful discussions; Jessica H. Hennacy for strains and reagents and for feedback on the manuscript; Eric Franklin for support with the TEMs and feedback on the figures; Angelo Kayser-Browne for feedback on the manuscript; and all present and former members of the Jonikas Lab for advice and discussions. Research reported in this publication was supported by the National Institute of General Medical Sciences (NIGMS) of the National Institutes of Health under grant numbers T32GM007388 and 1R01GM140032-01; by National Science Foundation Grant MCB-2410354; by the Bill and Melinda Gates Foundation and United Kingdom Foreign, Commonwealth & Development Office grant INV-054558; by the Howard Hughes Medical Institute; and by a grant to Princeton University from the Howard Hughes Medical Institute through the Gilliam Fellows Program. The content is solely the responsibility of the authors and does not necessarily represent the official views of the National Institutes of Health. We acknowledge the use of Princeton’s Imaging and Analysis Center (IAC), which is partially supported by the Princeton Center for Complex Materials (PCCM), a National Science Foundation Materials Research Science and Engineering Center (MRSEC; DMR-1420541). Confocal imaging was performed with support from the Confocal Imaging Facility, a Nikon Center of Excellence, in the Department of Molecular Biology at Princeton University. Molecular graphics and analyses of protein sequences were performed with UCSF ChimeraX, developed by the Resource for Biocomputing, Visualization, and Informatics at the University of California, San Francisco, with support from National Institutes of Health R01-GM129325 and the Office of Cyber Infrastructure and Computational Biology, National Institute of Allergy and Infectious Diseases.

## Author Contributions

V.L.C., M.I.B., and M.C.J. designed research; V.L.C. performed PCRs, growth assays, electron microscopy, crosses, iodine stain assays, and starch quantification assays; V.L.C. and M.I.B performed transformations; V.L.C. and A.G. performed confocal microscopy and western blots; M.I.B. performed cloning and starch binding assays; L.W. performed immunoprecipitation-mass spectrometry; V.L.C., M.I.B., A.G., and M.C.J. analyzed the data; V.L.C. and M.C.J wrote the paper; all authors contributed to editing the paper.

## References

1. L. C. M. Mackinder, et al., A repeat protein links Rubisco to form the eukaryotic carbon-concentrating organelle. Proc. Natl. Acad. Sci. U.S.A. 113, 5958–5963 (2016).

2. J. Barrett, P. Girr, L. C. M. Mackinder, Pyrenoids: CO2-fixing phase separated liquid organelles. Biochimica et Biophysica Acta (BBA) - Molecular Cell Research 1868, 118949 (2021).

3. S. He, V. L. Crans, M. C. Jonikas, The pyrenoid: the eukaryotic CO2-concentrating organelle. The Plant Cell 35, 3236–3259 (2023).

4. P. Flombaum, et al., Present and future global distributions of the marine Cyanobacteria Prochlorococcus and Synechococcus. Proc. Natl. Acad. Sci. U.S.A. 110, 9824–9829 (2013).

5. C. Rousseaux, W. Gregg, Interannual Variation in Phytoplankton Primary Production at A Global Scale. Remote Sensing 6, 1–19 (2013).

6. F. Not, et al., A Single Species, Micromonas pusilla (Prasinophyceae), Dominates the Eukaryotic Picoplankton in the Western English Channel. Appl Environ Microbiol 70, 4064– 4072 (2004).

7. H. R. Thierstein, J. R. Young, Eds., Coccolithophores (Springer Berlin Heidelberg, 2004).

8. M. Meyer, H. Griffiths, Origins and diversity of eukaryotic CO2-concentrating mechanisms: lessons for the future. Journal of Experimental Botany 64, 769–786 (2013).

9. B. D. Rae, et al., Progress and challenges of engineering a biophysical CO2-concentrating mechanism into higher plants. Journal of Experimental Botany 68, 3717–3737 (2017).

10. J. H. Hennacy, M. C. Jonikas, Prospects for Engineering Biophysical CO _2_ Concentrating Mechanisms into Land Plants to Enhance Yields. Annu. Rev. Plant Biol. 71, 461–485 (2020).

11. L. Adler, et al., New horizons for building pyrenoid-based CO2-concentrating mechanisms in plants to improve yields. Plant Physiology 190, 1609–1627 (2022).

12. A. A. Hyman, C. A. Weber, F. Jülicher, Liquid-Liquid Phase Separation in Biology. Annu. Rev. Cell Dev. Biol. 30, 39–58 (2014).

13. D. M. Mitrea, R. W. Kriwacki, Phase separation in biology; functional organization of a higher order. Cell Commun Signal 14, 1 (2016).

14. S. F. Banani, H. O. Lee, A. A. Hyman, M. K. Rosen, Biomolecular condensates: organizers of cellular biochemistry. Nat Rev Mol Cell Biol 18, 285–298 (2017).

15. Y. Shin, C. P. Brangwynne, Liquid phase condensation in cell physiology and disease. Science 357, eaaf4382 (2017).

16. E. S. Freeman Rosenzweig, et al., The Eukaryotic CO2-Concentrating Organelle Is Liquid-like and Exhibits Dynamic Reorganization. Cell 171, 148-162.e19 (2017).

17. B. D. Engel, et al., Native architecture of the Chlamydomonas chloroplast revealed by in situ cryo-electron tomography. eLife 4, e04889 (2015).

18. J. H. Hennacy, et al., SAGA1 and MITH1 produce matrix-traversing membranes in the CO_2_-fixing pyrenoid. Nat. Plants 10, 2038–2051 (2024).

19. R. Sager, G. E. Palade, Structure and Development of the Chloroplast in Chlamydomonas. The Journal of Cell Biology 3, 463–488 (1957).

20. A. Bracher, S. M. Whitney, F. U. Hartl, M. Hayer-Hartl, Biogenesis and Metabolic Maintenance of Rubisco. Annu. Rev. Plant Biol. 68, 29–60 (2017).

21. N. A. Pronina, S. Avramova, D. Georgiev, V. E. Semenenko, A pattern of carbonic anhydrase activity in Chlorella and Scenedesmus on cell adaptation to high level light intensity and low CO2 concentration. Fiziologia Rastenii 28, 43–52 (1981).

22. J. A. Raven, CO _2_-concentrating mechanisms: a direct role for thylakoid lumen acidification? Plant Cell & Environment 20, 147–154 (1997).

23. C. Fei, A. T. Wilson, N. M. Mangan, N. S. Wingreen, M. C. Jonikas, Modelling the pyrenoid-based CO2-concentrating mechanism provides insights into its operating principles and a roadmap for its engineering into crops. Nat. Plants 8, 583–595 (2022).

24. C. Toyokawa, T. Yamano, H. Fukuzawa, Pyrenoid Starch Sheath Is Required for LCIB Localization and the CO _2_-Concentrating Mechanism in Green Algae. Plant Physiol. 182, 1883–1893 (2020).

25. P. Forssell, Oxygen permeability of amylose and amylopectin films. Carbohydrate Polymers 47, 125–129 (2002).

26. I. Arvanitoyannis, M. Kalichevsky, J. M. V. Blanshard, E. Psomiadou, Study of diffusion and permeation of gases in undrawn and uniaxially drawn films made from potato and rice starch conditioned at different relative humidities. Carbohydrate Polymers 24, 1–15 (1994).

27. M. T. Meyer, et al., Assembly of the algal CO _2_-fixing organelle, the pyrenoid, is guided by a Rubisco-binding motif. Sci. Adv. 6, eabd2408 (2020).

28. S. He, et al., The structural basis of Rubisco phase separation in the pyrenoid. Nat. Plants 6, 1480–1490 (2020).

29. A. K. Itakura, et al., A Rubisco-binding protein is required for normal pyrenoid number and starch sheath morphology in Chlamydomonas reinhardtii. Proc. Natl. Acad. Sci. U.S.A. 116, 18445–18454 (2019).

30. R. Zhang, et al., High-Throughput Genotyping of Green Algal Mutants Reveals Random Distribution of Mutagenic Insertion Sites and Endonucleolytic Cleavage of Transforming DNA. Plant Cell 26, 1398–1409 (2014).

31. N. Atkinson, R. Stringer, S. R. Mitchell, D. Seung, A. J. McCormick, SAGA1 and SAGA2 promote starch formation around proto-pyrenoids in Arabidopsis chloroplasts. Proc. Natl. Acad. Sci. U.S.A. 121, e2311013121 (2024).

32. X. Li, et al., A genome-wide algal mutant library and functional screen identifies genes required for eukaryotic photosynthesis. Nat Genet 51, 627–635 (2019).

33. K. Kuchitsu, M. Tsuzuki, S. Miyachi, Changes of Starch Localization within the Chloroplast Induced by Changes in CO_2_ Concentration during Growth of Chlamydomonas reinhardtii: Independent Regulation of Pyrenoid Starch and Stroma Starch. Plant and Cell Physiology (1988). 10.1093/oxfordjournals.pcp.a077635.

34. K. Kuchitsu, M. Tsuzuki, S. Miyachi, Polypeptide composition and enzyme activities of the pyrenoid and its regulation by CO _2_ concentration in unicellular green algae. Can. J. Bot. 69, 1062–1069 (1991).

35. Z. Ramazanov, et al., The induction of the CO2-concentrating mechanism is correlated with the formation of the starch sheath around the pyrenoid of Chlamydomonas reinhardtii. Planta 195 (1994).

36. T. Yamano, et al., Light and Low-CO2-Dependent LCIB–LCIC Complex Localization in the Chloroplast Supports the Carbon-Concentrating Mechanism in Chlamydomonas reinhardtii. Plant and Cell Physiology 51, 1453–1468 (2010).

37. S. Ichikawa, M. Sakata, T. Oba, Y. Kodama, Fluorescein staining of chloroplast starch granules in living plants. Plant Physiol 194, 662–672 (2024).

38. C. Zabawinski, et al., Starchless Mutants of Chlamydomonas reinhardtii Lack the Small Subunit of a Heterotetrameric ADP-Glucose Pyrophosphorylase. J Bacteriol 183, 1069– 1077 (2001).

39. H. Brust, S. Orzechowski, J. Fettke, Starch and Glycogen Analyses: Methods and Techniques. Biomolecules 10, 1020 (2020).

40. L. C. M. Mackinder, et al., A Spatial Interactome Reveals the Protein Organization of the Algal CO2-Concentrating Mechanism. Cell 171, 133-147.e14 (2017).

41. N. J. Belshaw, G. Williamson, Interaction of β-cyclodextrin with the granular starch binding domain of glucoamylase. Biochimica et Biophysica Acta (BBA) - Protein Structure and Molecular Enzymology 1078, 117–120 (1991).

42. N. J. Belshaw, G. Williamson, Specificity of the binding domain of glucoamylase 1. European Journal of Biochemistry 211, 717–724 (1993).

43. D. Seung, et al., Homologs of PROTEIN TARGETING TO STARCH Control Starch Granule Initiation in Arabidopsis Leaves. Plant Cell 29, 1657–1677 (2017).

44. S. Wüpper, K. Lüersen, G. Rimbach, Cyclodextrins, Natural Compounds, and Plant Bioactives—A Nutritional Perspective. Biomolecules 11, 401 (2021).

45. M. R. Abt, S. C. Zeeman, Evolutionary innovations in starch metabolism. Current Opinion in Plant Biology 55, 109–117 (2020).

46. D. Seung, T. B. Schreier, L. Bürgy, S. Eicke, S. C. Zeeman, Two Plastidial Coiled-Coil Proteins Are Essential for Normal Starch Granule Initiation in Arabidopsis. Plant Cell 30, 1523–1542 (2018).

47. A. Courseaux, P. Deschamps, D. Dauvillée, An alternative pathway to starch granule initiation unraveled in Chlamydomonas reinhardtii. Plant And Cell Physiology 66, 738–752 (2025).

48. A. J. M. Heutinck, et al., Branched oligosaccharides cause atypical starch granule initiation in Arabidopsis chloroplasts. Plant Physiology 197, kiaf002 (2025).

49. J. Raven, Contributions of anoxygenic and oxygenic phototrophy and chemolithotrophy to carbon and oxygen fluxes in aquatic environments. Aquat. Microb. Ecol. 56, 177–192 (2009).

50. J. C. Villarreal, S. S. Renner, Hornwort pyrenoids, carbon-concentrating structures, evolved and were lost at least five times during the last 100 million years. Proc. Natl. Acad. Sci. U.S.A. 109, 18873–18878 (2012).

51. H. Griffiths, M. T. Meyer, R. E. M. Rickaby, Overcoming adversity through diversity: aquatic carbon concentrating mechanisms. Journal of Experimental Botany 68, 3689–3695 (2017).

52. J. Barrett, et al., A promiscuous mechanism to phase separate eukaryotic carbon fixation in the green lineage. Nat. Plants 10, 1801–1813 (2024).

53. Z. G. Oh, et al., A linker protein from a red-type pyrenoid phase separates with Rubisco via oligomerizing sticker motifs. Proc. Natl. Acad. Sci. U.S.A. 120, e2304833120 (2023).

54. J. Kropat, et al., A revised mineral nutrient supplement increases biomass and growth rate in Chlamydomonas reinhardtii. The Plant Journal 66, 770–780 (2011).

55. X. Jiang, D. Stern, Mating and Tetrad Separation of Chlamydomonas reinhardtii for Genetic Analysis. JoVE 1274 (2009). 10.3791/1274.

56. J. Schindelin, et al., Fiji: an open-source platform for biological-image analysis. Nat Methods 9, 676–682 (2012).

57. L. Wang, et al., A chloroplast protein atlas reveals punctate structures and spatial organization of biosynthetic pathways. Cell 186, 3499-3518.e14 (2023).

58. B. Delrue, et al., Waxy Chlamydomonas reinhardtii: monocellular algal mutants defective in amylose biosynthesis and granule-bound starch synthase activity accumulate a structurally modified amylopectin. J Bacteriol 174, 3612–3620 (1992).

59. R. J. Craig, et al., The Chlamydomonas Genome Project, version 6: Reference assemblies for mating-type plus and minus strains reveal extensive structural mutation in the laboratory. The Plant Cell 35, 644–672 (2023).

60. M. Blum, et al., InterPro: the protein sequence classification resource in 2025. Nucleic Acids Research 53, D444–D456 (2025).

61. K. Sorimachi, M.-F.L. Gal-Coëffet, G. Williamson, D. B. Archer, M. P. Williamson, Solution structure of the granular starch binding domain of Aspergillus niger glucoamylase bound to β-cyclodextrin. Structure 5, 647–661 (1997).

62. M. Goujon, et al., A new bioinformatics analysis tools framework at EMBL-EBI. Nucleic Acids Research 38, W695–W699 (2010).

63. F. Sievers, et al., Fast, scalable generation of high-quality protein multiple sequence alignments using Clustal Omega. Molecular Systems Biology 7, 539 (2011).

64. J. Abramson, et al., Accurate structure prediction of biomolecular interactions with AlphaFold 3. Nature 630, 493–500 (2024).

65. K. Sorimachi, M.-F. Le Gal-Coeffet, G. Williamson, D. B. Archer, M. P. Williamson, GLUCOAMYLASE, GRANULAR STARCH-BINDING DOMAIN COMPLEX WITH CYCLODEXTRIN, NMR, 5 STRUCTURES: 1acz. 10.2210/pdb1acz/pdb. Deposited 7 July 1997.

66. K. Sorimachi, et al., GLUCOAMYLASE, GRANULAR STARCH-BINDING DOMAIN, NMR, 5 STRUCTURES: 1kul. 10.2210/pdb1kul/pdb. Deposited 11 July 1996.

67. K. Sorimachi, et al., Solution Structure of the Granular Starch Binding Domain of Glucoamylase fromAspergillus nigerby Nuclear Magnetic Resonance Spectroscopy. Journal of Molecular Biology 259, 970–987 (1996).

68. H. M. Berman, et al., The Protein Data Bank. Nucleic Acids Research 28, 235–242 (2000).

69. S. Bittrich, J. Segura, J. M. Duarte, S. K. Burley, Y. Rose, RCSB protein Data Bank: exploring protein 3D similarities via comprehensive structural alignments. Bioinformatics 40, btae370 (2024).

70. E. C. Meng, et al., UCSF ChimeraX : Tools for structure building and analysis. Protein Science 32, e4792 (2023).

