## Supporting Information for "SAGA1 and SAGA2 localize the starch sheath to the pyrenoid in *Chlamydomonas reinhardtii*"

**This PDF file includes:**

Figures S1 to S18
Tables S1 to S5
SI Materials and Methods
SI References

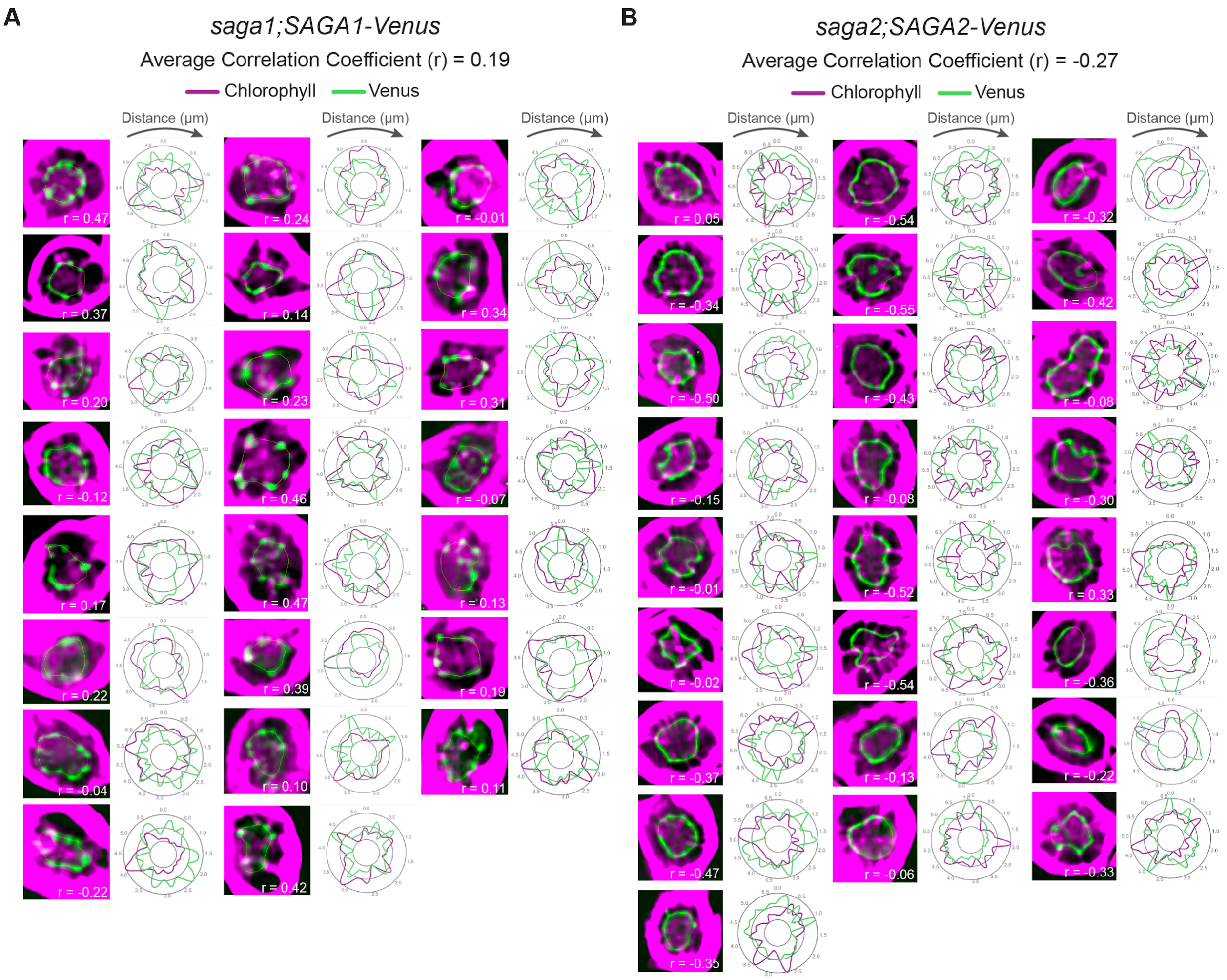

**Figure S1**. Confocal images of *saga1;SAGA1-Venus* (A) and *saga2;SAGA2-Venus* (B). Graphs to the right of each image display quantifications of normalized fluorescence values for SAGA1-Venus, SAGA2-Venus, and chlorophyll autofluorescence around the periphery of the pyrenoids. Yellow circles indicate the lines that were used to measure fluorescence intensities. Pearson's correlation coefficients (r) are labeled on the confocal images.

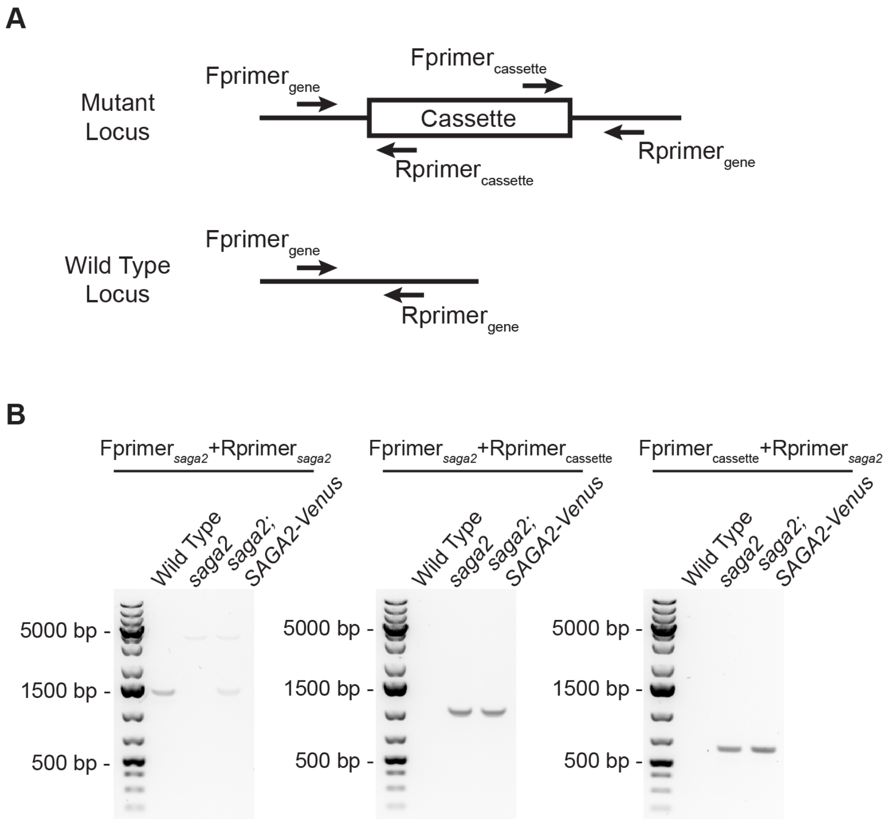

**Figure S2.** The *saga2* mutant contains the insertional cassette. (A) Schematic of the PCR confirmation strategy for CLiP library mutants. (B) The Fprimer$_{saga2}$/Rprimer$_{saga2}$ primer pair yields a ~1,500 bp band in wild type and a faint ~4,000 bp band in *saga2* due to the presence of the insertional cassette. The *saga2;SAGA2-Venus* lane has both of these bands due to the presence of both the *saga2* insertional cassette and the re-introduced wild type *SAGA2* gene. The Fprimer$_{saga2}$/Rprimer$_{cassette}$ pair yields no band in wild type and a ~1,000 bp band in *saga2* and *saga2;SAGA2-Venus*, and the Fprimer$_{cassette}$/Rprimer$_{saga2}$ primer pair yields no band in wild type and a ~600 bp band in *saga2* and *saga2;SAGA2-Venus*.

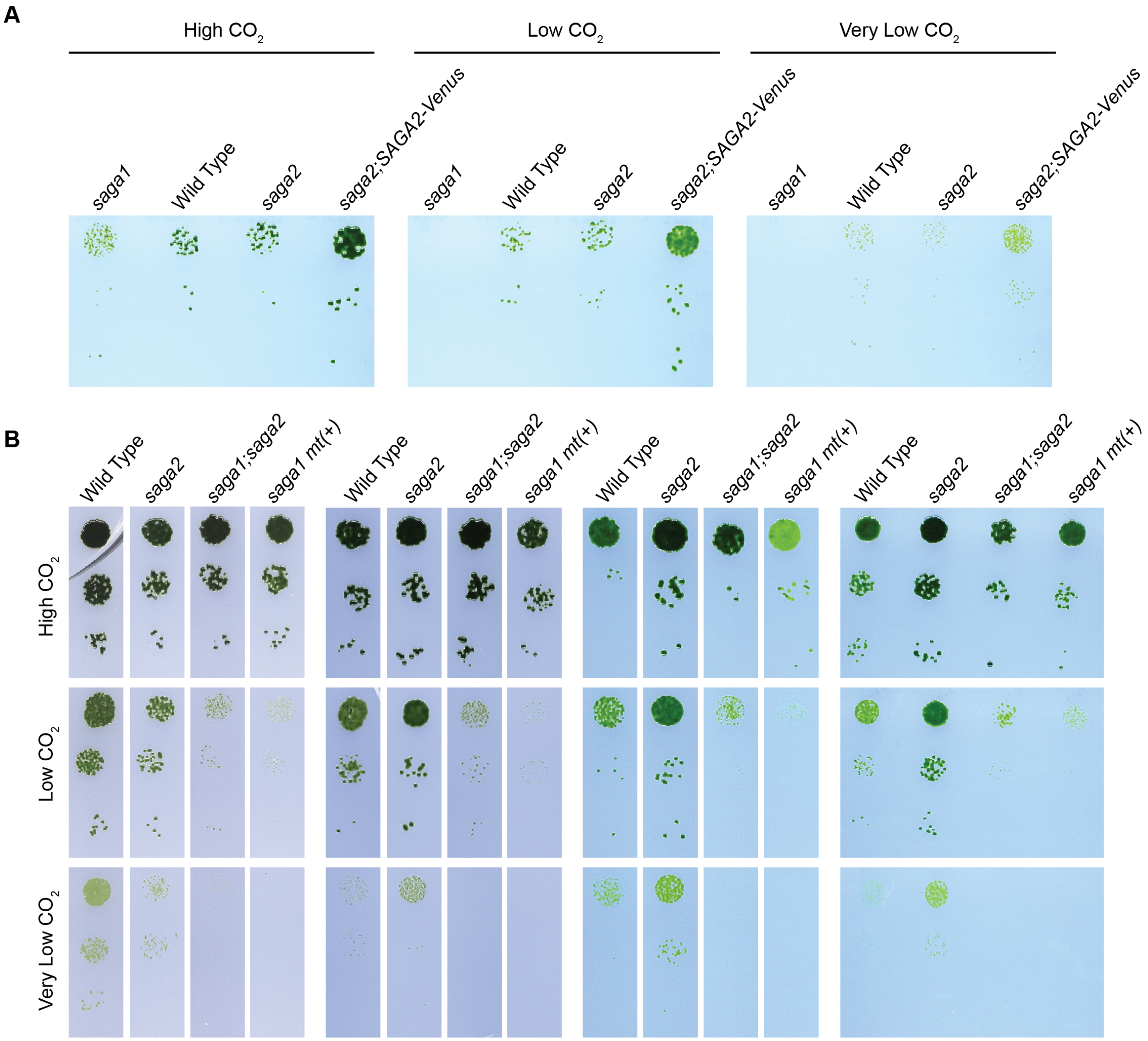

**Figure S3**. Agar spot test growth phenotypes of wild type, *saga1*, *saga2*, and *saga2;SAGA2-Venus* (A), and wild type, *saga1 mt(+)*, *saga2*, and *saga1;saga2* (B). Liquid cultures were spotted on TP minimal medium and grown at high (3% v/v), low (0.04% v/v), and very low (<0.004% v/v) $CO_2$ under 100 µmol photons·$m^{-2}$·$s^{-1}$ illumination for 7 days.

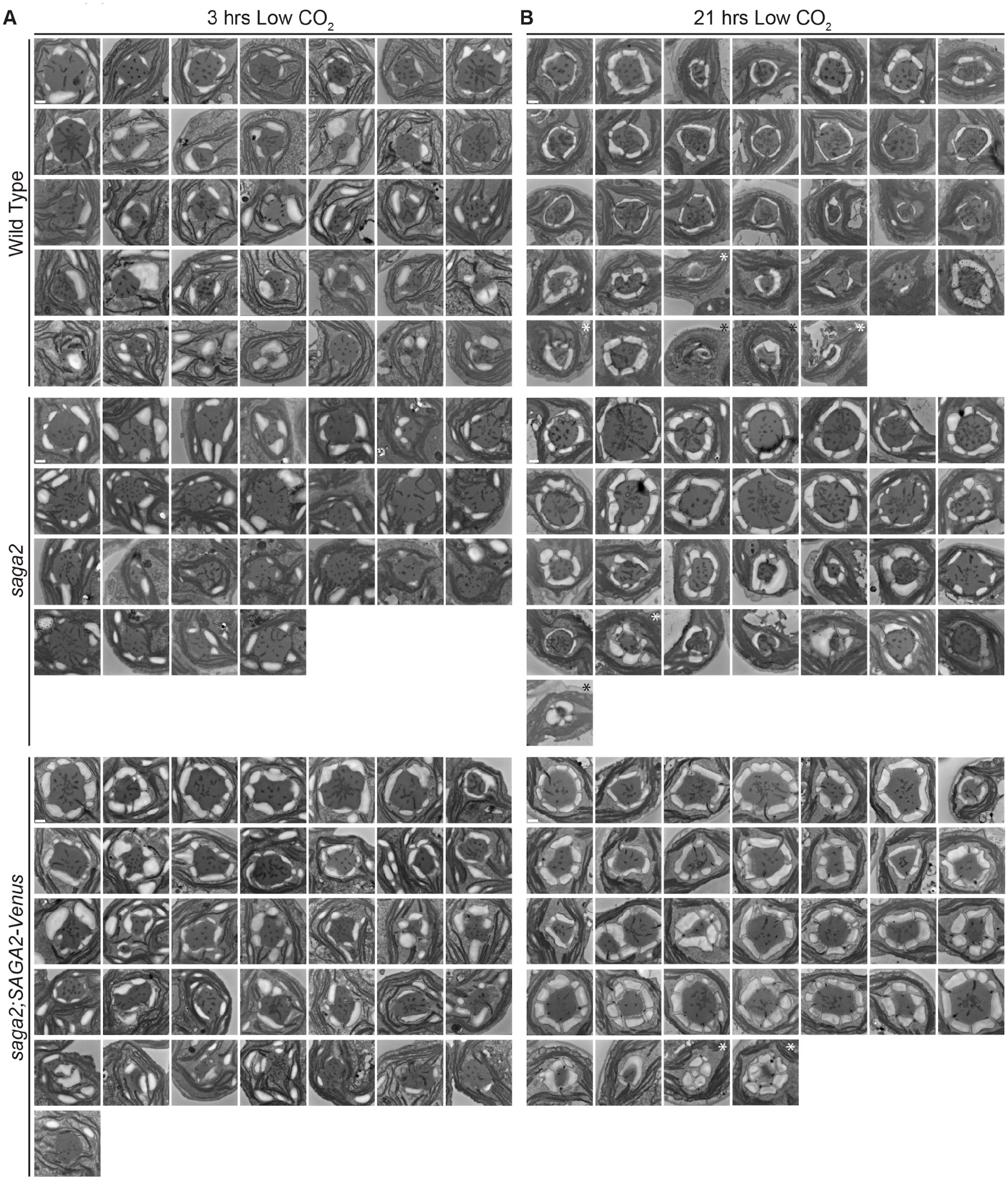

**Figure S4.** TEM images of pyrenoids in wild type, *saga2*, and *saga2;SAGA2-Venus*. (A) Images of pyrenoids from cells grown at high (3% v/v) $CO_2$ in minimal media, then moved to low (0.04% v/v) $CO_2$ for 3 hours before harvesting. (B) Images of pyrenoids from cells grown at high (3% v/v) $CO_2$ in minimal media, then moved to low (0.04% v/v) $CO_2$ for 21 hours before harvesting. Scale bars = 500 nm. Asterisks mark abnormal pyrenoids that were excluded from the quantifications in Fig. 2 *C* and *E* and Fig. S5.

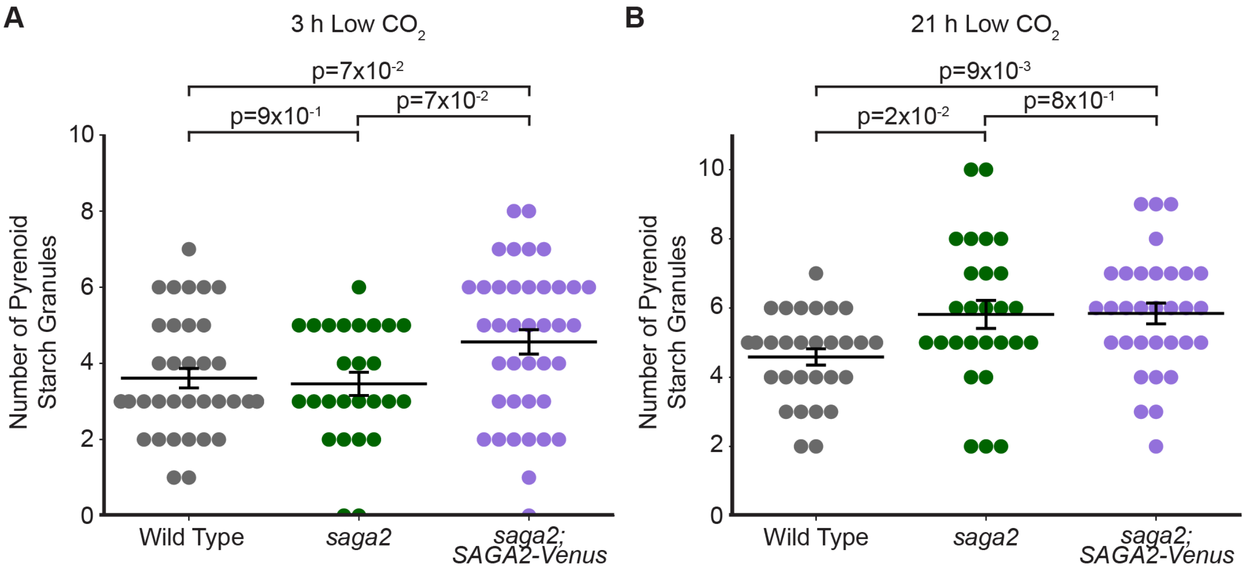

**Figure S5**. Number of starch granules in contact with pyrenoids in wild type, *saga2*, and *saga2;SAGA2-Venus* TEMs (Fig. 2 *B* and *D*, Fig. S4). (A) Quantifications of the number of pyrenoid starch granules in cells grown at high (3% v/v) $CO_2$ in minimal media, then moved to low (0.04% v/v) $CO_2$ for 3 hours before harvesting. (B) Quantifications of the number of pyrenoid starch granules in cells grown at high (3% v/v) $CO_2$ in minimal media, then moved to low (0.04% v/v) $CO_2$ for 21 hours before harvesting. Statistical p-values were calculated using Kruskal-Wallis ($p=3\times10^{-2}$ in A, $p=5\times10^{-3}$ in B) followed by Dunn's multiple comparisons test. Error bars represent the standard error of the mean.

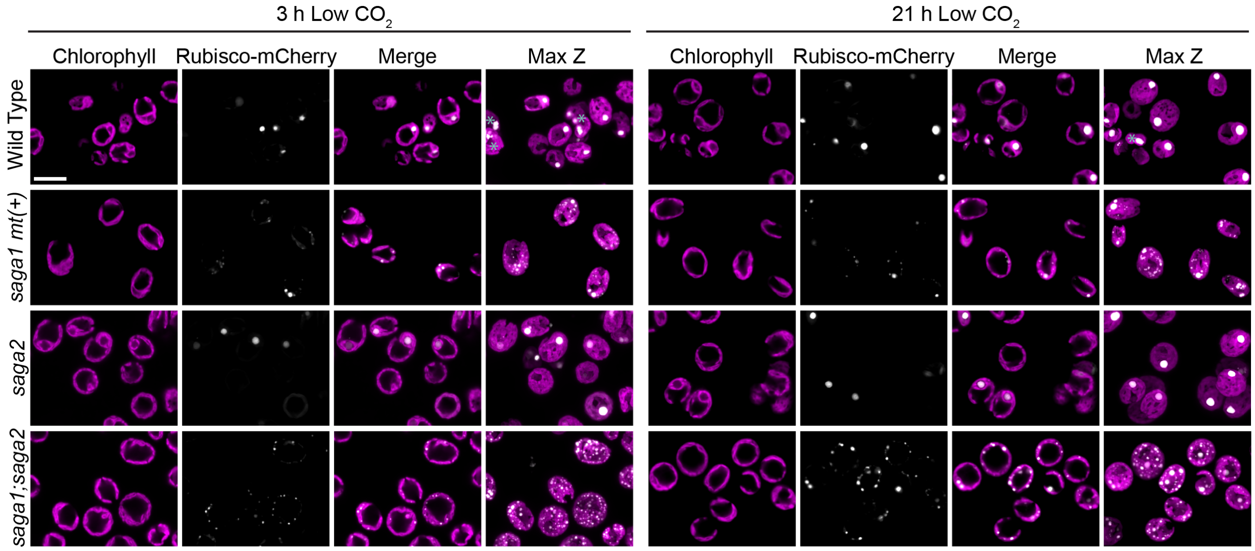

**Figure S6.** Localization of RBCS1-mCherry in wild type and *saga* mutant backgrounds. Cells were grown at high (3% v/v) $CO_2$ in minimal media, then moved to low (0.04% v/v) $CO_2$ for 3 hours (left) and 21 hours (right) before imaging. Scale bar = 10 µm. Asterisks mark cells that moved during the process of acquiring Z-stacks, causing them to blur in the Max Z projections.

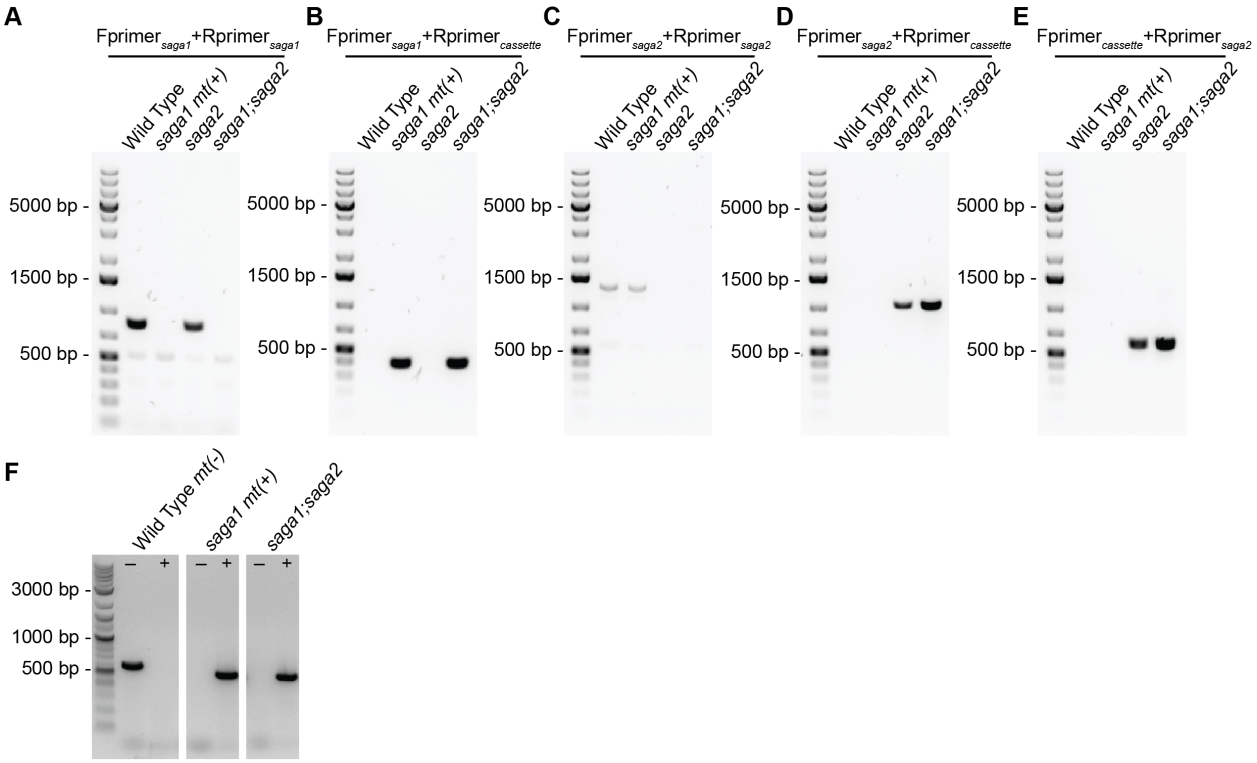

**Figure S7.** The *saga1;saga2* double mutant contains insertional cassettes in both the *saga1* and the *saga2* loci, does not express the SAGA1 protein, and is mating type plus (*mt(+)*). See Fig. S2*A* for primer schematic. (A) The Fprimer$_{saga1}$/Rprimer$_{saga1}$ primer pair yields a ~800 bp band in wild type and *saga2* but no band in *saga1 mt(+)* and *saga1;saga2*. (B) The Fprimer$_{saga1}$/Rprimer$_{cassette}$ primer pair yields a ~400 bp band in *saga1 mt(+)* and *saga1;saga2* but no band in wild type and *saga2*. (C) The Fprimer$_{saga2}$/Rprimer$_{saga2}$ primer pair yields a ~1,400 bp band in wild type and *saga1 mt(+)* but no band in *saga2* and *saga1;saga2*. (D) The Fprimer$_{saga2}$/Rprimer$_{cassette}$ primer pair yields a ~1,000 bp band in *saga2* and *saga1;saga2* but no band in wild type and *saga1 mt(+)*. (E) The Fprimer$_{cassette}$/Rprimer$_{saga2}$ primer pair yields a ~600 bp band in *saga2* and *saga1;saga2* but no band in wild type and *saga1 mt(+)*. (F) PCR results checking the mating type (*mt*) locus of wild type (which is *mt(-)*), *saga1 mt(+)*, and *saga1;saga2*. Left lanes are primers that bind to the *mt(-)* locus and right lanes are primers that bind to the *mt(+)* locus. The *mt(-)* primers yield a ~600 bp band in wild type but no band in *saga1 mt(+)* and *saga1;saga2*. The *mt(+)* primers yield a ~400 bp band in *saga1 mt(+)* and *saga1;saga2* but no band in wild type.

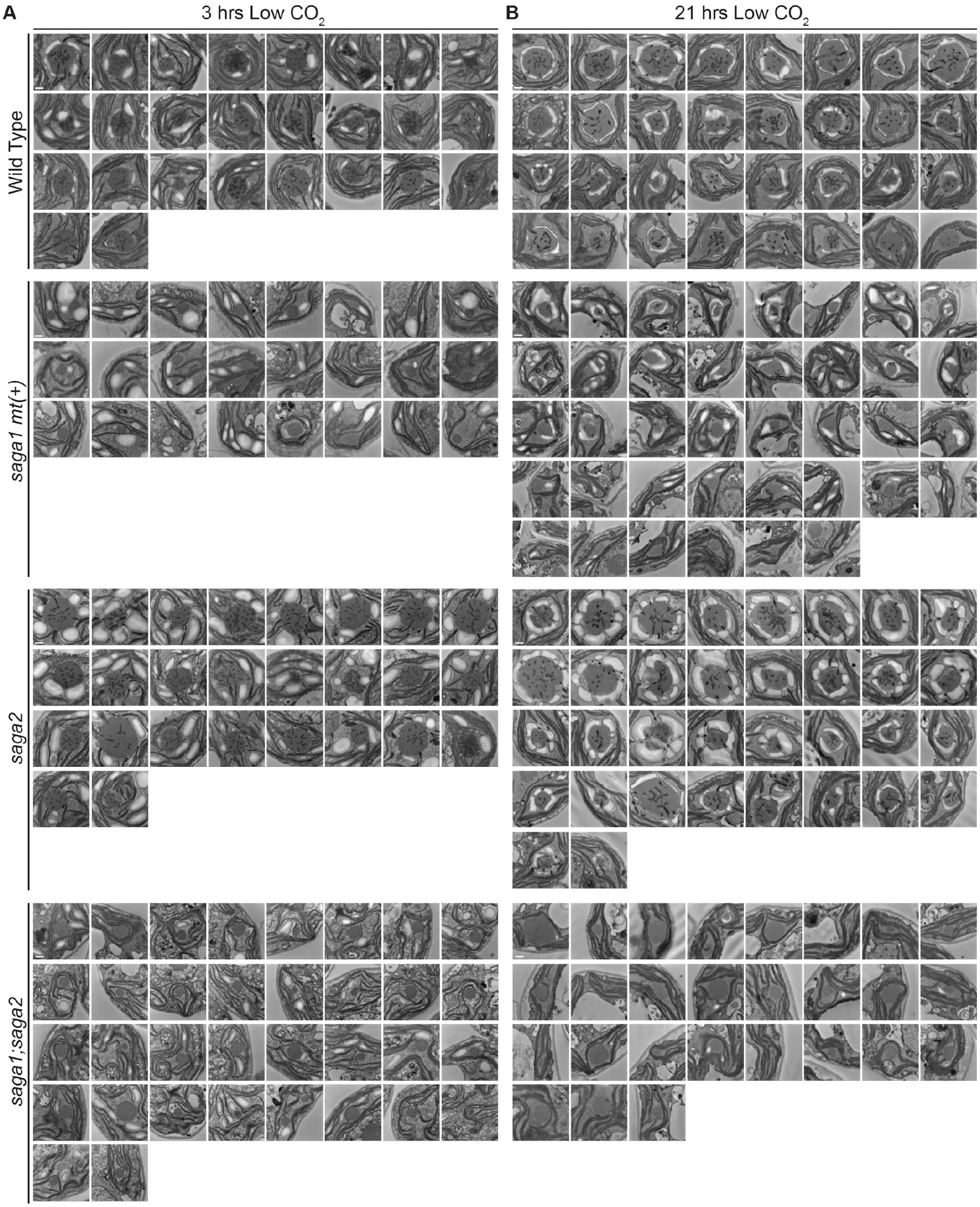

**Figure S8.** TEM images of pyrenoids in wild type, *saga1 mt(+)*, *saga2*, and *saga1;saga2*. (A) TEM images of pyrenoids in cells that were grown at high (3% v/v) $CO_2$ in minimal media, then moved to low (0.04% v/v)

$CO_2$ for 3 hours before harvesting. (B) TEM images of pyrenoids in cells that were grown at high (3% v/v) $CO_2$ in minimal media, then moved to low (0.04% v/v) $CO_2$ for 21 hours before harvesting. Scale bars = 500 nm.

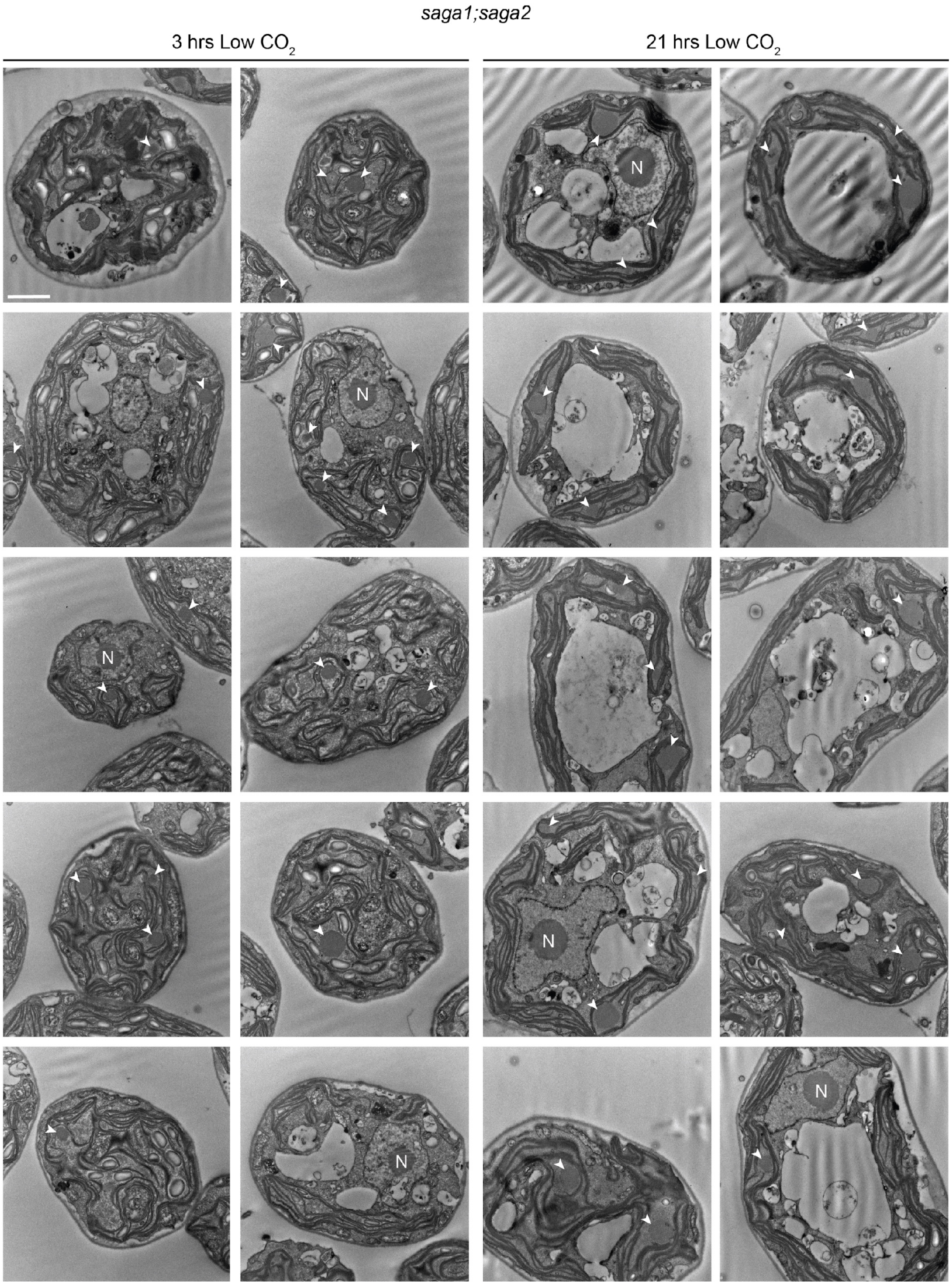

**Figure S9.** Whole cell TEM images of the *saga1;saga2* double mutant. Cells were grown at high (3% v/v) $CO_2$ in minimal media, then moved to low (0.04% v/v) $CO_2$ for 3 hours (left) and 21 hours (right) before imaging. Arrows indicate pyrenoids. N: Nucleolus. Scale bar = 2 μm.

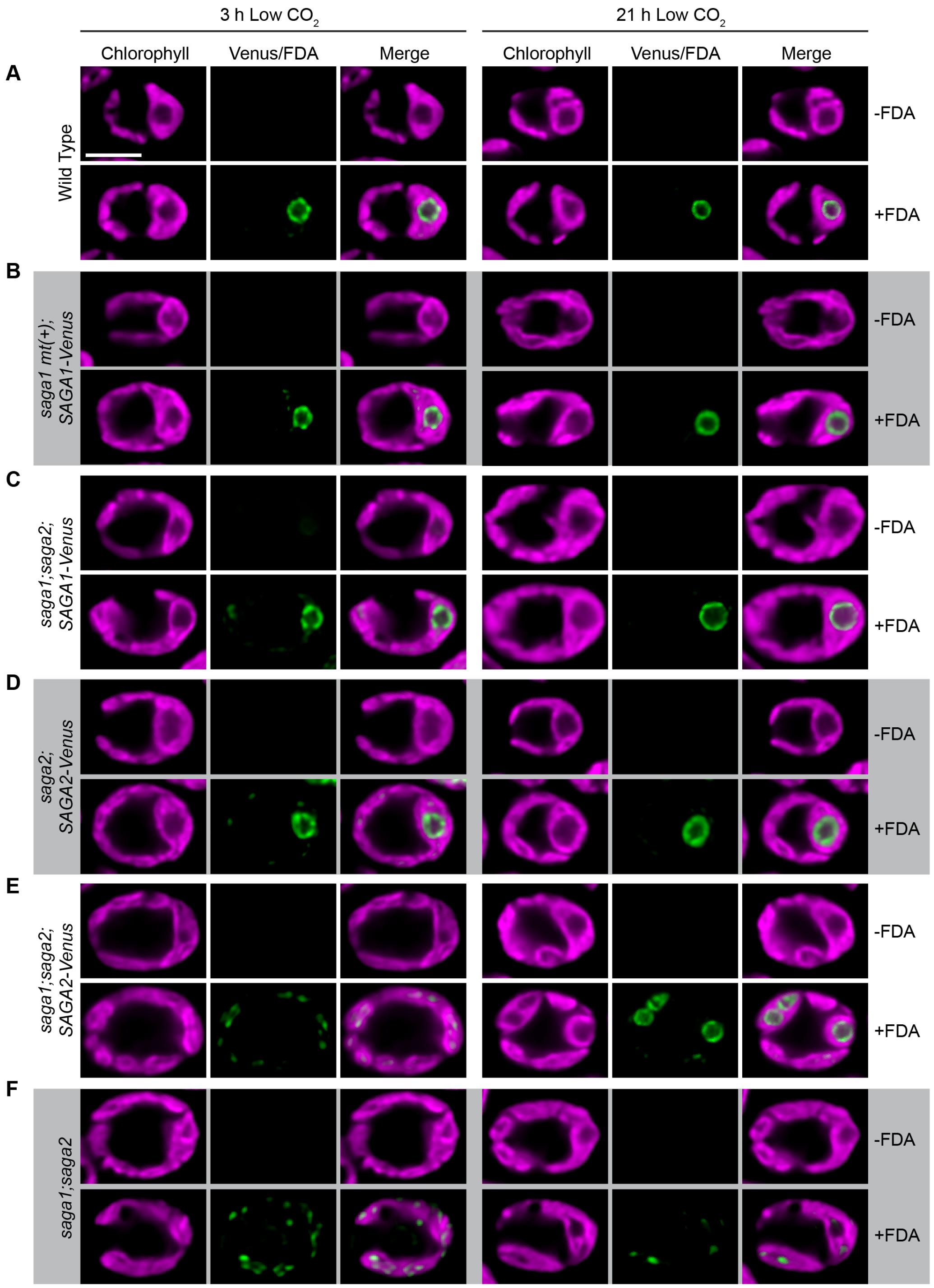

**Figure S10**. Localization of Fluorescein (FDA) in wild type (A), *saga1 mt(+);SAGA1-Venus* (B), *saga1;saga2;SAGA1-Venus* (C), *saga2;SAGA2-Venus* (D), *saga1;saga2;SAGA2-Venus* (E), and *saga1;saga2* (F). Cells were grown at high (3% v/v) $CO_2$ in minimal media, then moved to low (0.04% v/v) $CO_2$ for 3 hours (left) and 21 hours (right) before imaging. For each strain the top row shows cells that were not stained with FDA, with the Venus/FDA channel set to the same display intensity as the image in the row below of a cell that was stained with FDA. Although both fluorescein and Venus have similar excitation and emission spectra, the fluorescein signal was bright enough that the Venus signal was not visible when display values were set so that the fluorescein signal was not oversaturated, allowing us to visualize fluorescein despite the presence of Venus. Scale bar = 5 μm.

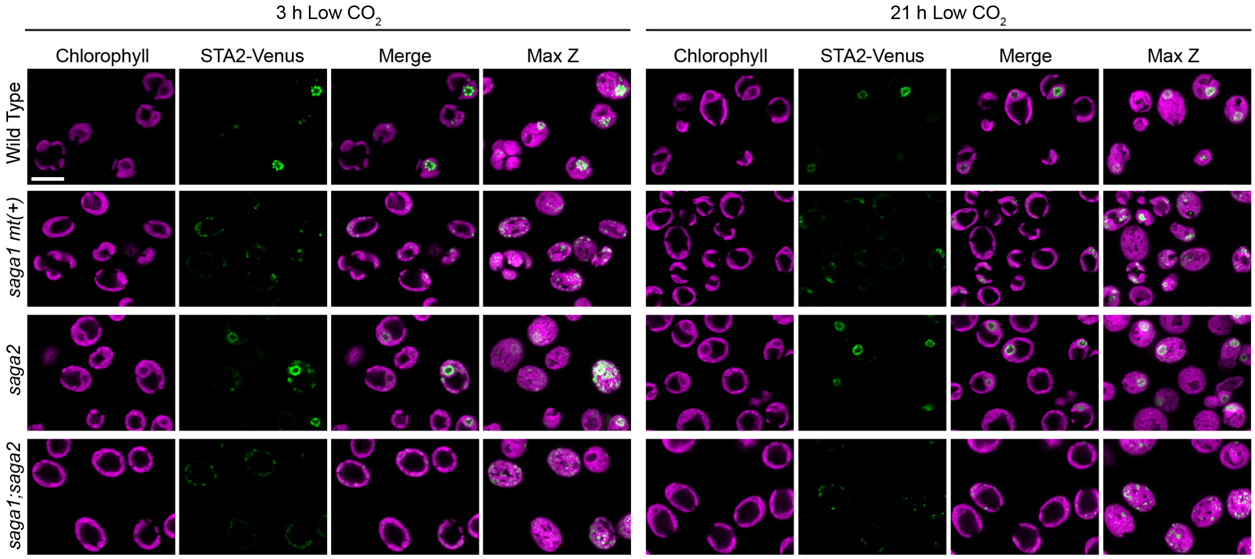

**Figure S11.** Localization of STA2-Venus in wild type and *saga* mutant backgrounds. Cells were grown at high (3% v/v) $CO_2$ in minimal media, then moved to low (0.04% v/v) $CO_2$ for 3 hours (left) and 21 hours (right) before imaging. Scale bar = 10 µm.

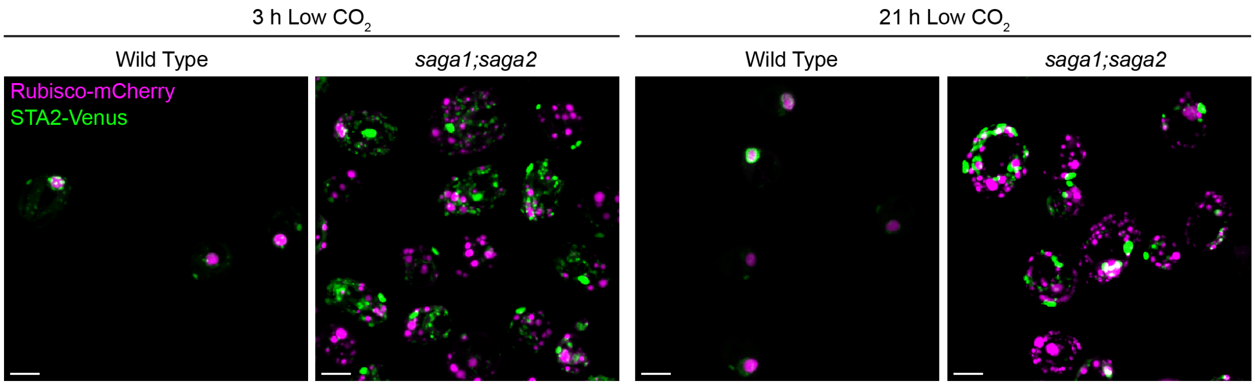

**Figure S12**. Max Z projections of Rubisco-mCherry and STA2-Venus in wild type and *saga1;saga2*. Cells were grown at high (3% v/v) $CO_2$ in minimal media, then moved to low (0.04% v/v) $CO_2$ for 3 hours (left) and 21 hours (right) before imaging.  Scale bars = 5 µm.

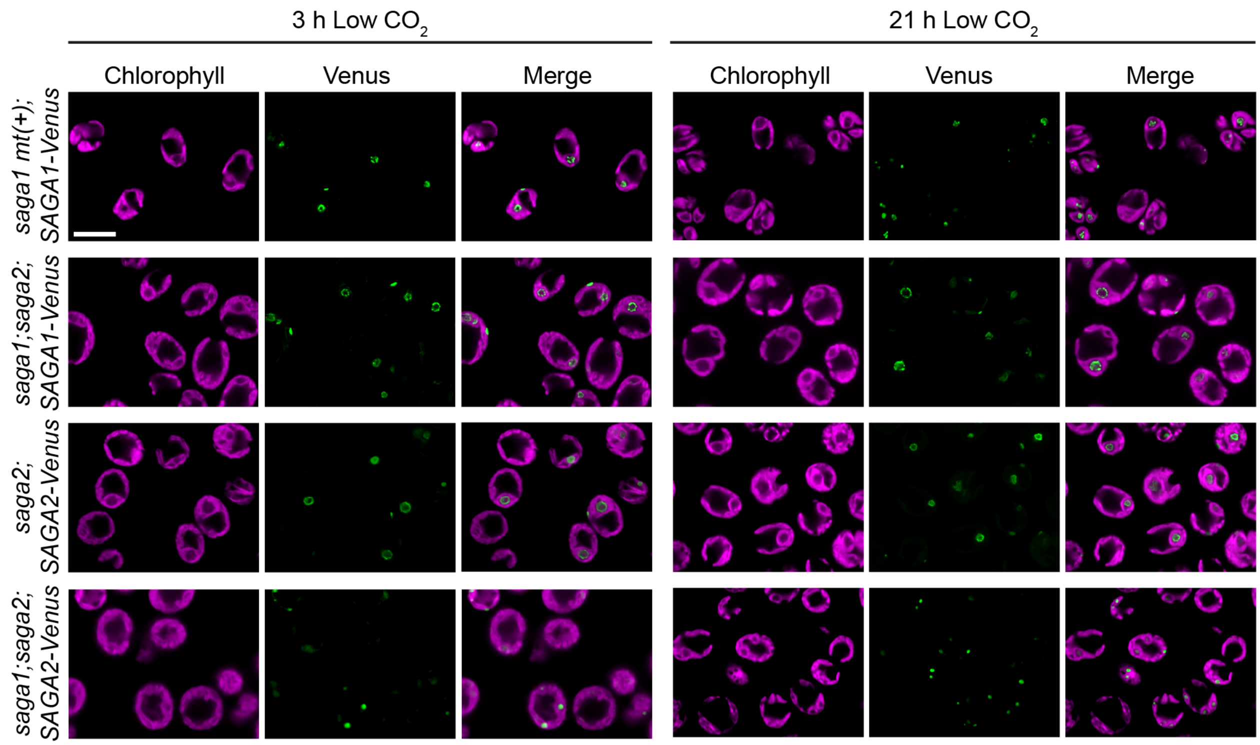

**Figure S13**. Localization of SAGA1-Venus in *saga1 mt(+)* and *saga1;saga2* and SAGA2-Venus in *saga2* and *saga1;saga2*. Cells were grown at high (3% v/v) $CO_2$ in minimal media, then moved to low (0.04% v/v) $CO_2$ for 3 hours (left) and 21 hours (right) before imaging. Scale bar = 10 µm.

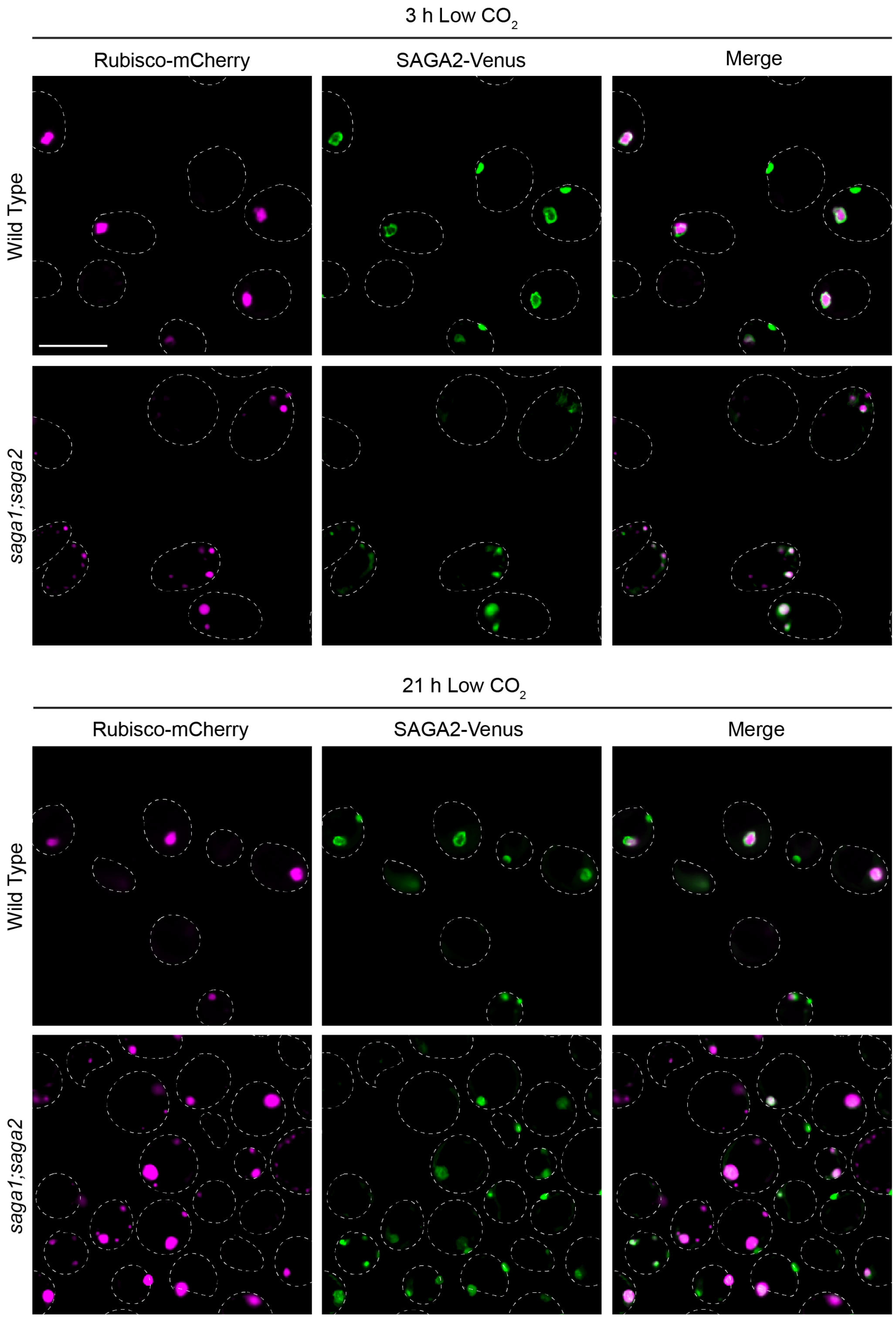

**Figure S14**. Localization of SAGA2-Venus and Rubisco-mCherry in wild type and *saga1;saga2*. Cells were grown at high (3% v/v) $CO_2$ in minimal media, then moved to low (0.04% v/v) $CO_2$ for 3 hours (top) and 21

hours (bottom) before imaging. Dotted lines outline cells based on chlorophyll autofluorescence (not shown). Scale bar = 10 μm.

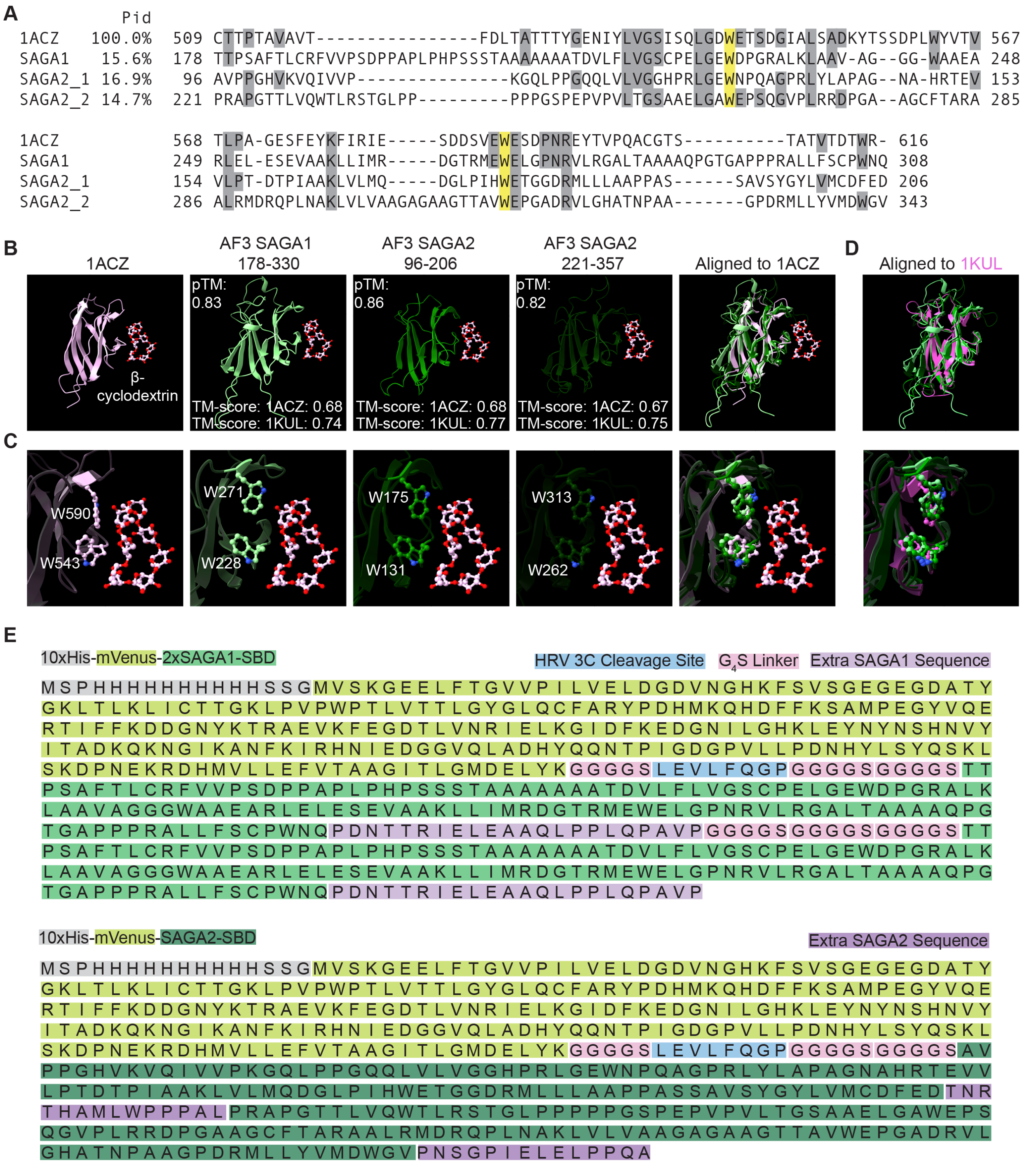

**Figure S15**. Sequence and structural analysis of the predicted SAGA1 and SAGA2 CBM20 domains. (A) Clustal Omega multiple sequence alignment of the predicted CBM20 sequences from SAGA1 and SAGA2 against the *Aspergillus niger* glucoamylase G1 CBM20 sequence from PDB (1ACZ). Grey letters indicate overall sequence homology to 1ACZ, while yellow letters indicate the location of the conserved tryptophan residues required for starch binding. Pid – percent identity. (B) RCSB PDB pairwise alignment of the

AlphaFold 3 (AF3) predicted structures of the SAGA1 and SAGA2 predicted CBM20 domains compared to the NMR solution structure of the *A. niger* glucoamylase G1 CBM20 in complex with β-cyclodextrin at the starch-binding site (1ACZ). For each SAGA CBM20 structure, β-cyclodextrin is shown based on its position relative to the SAGA CBM20's in the alignment with 1ACZ. (C) Close-up of the conserved tryptophan residues mediating starch binding in the structures from B. (D) Pairwise structural alignment of the SAGA1 and SAGA2 predicted CBM20 domains against the NMR solution structure of the free *A. niger* glucoamylase G1 CBM20 without β-cyclodextrin (1KUL). (E) Amino acid sequences of 2xSAGA1-SBD (top) and SAGA2-SBD (bottom). Gray: 10xHis tag. Yellow-green: mVenus. Green: SAGA1 CBM20. Blue: HRV 3C cleavage site. Pink: $G_4S$ linker. Light purple: extra native SAGA1 sequence. Dark green: SAGA2 CBM20. Dark purple: native SAGA2 sequence.

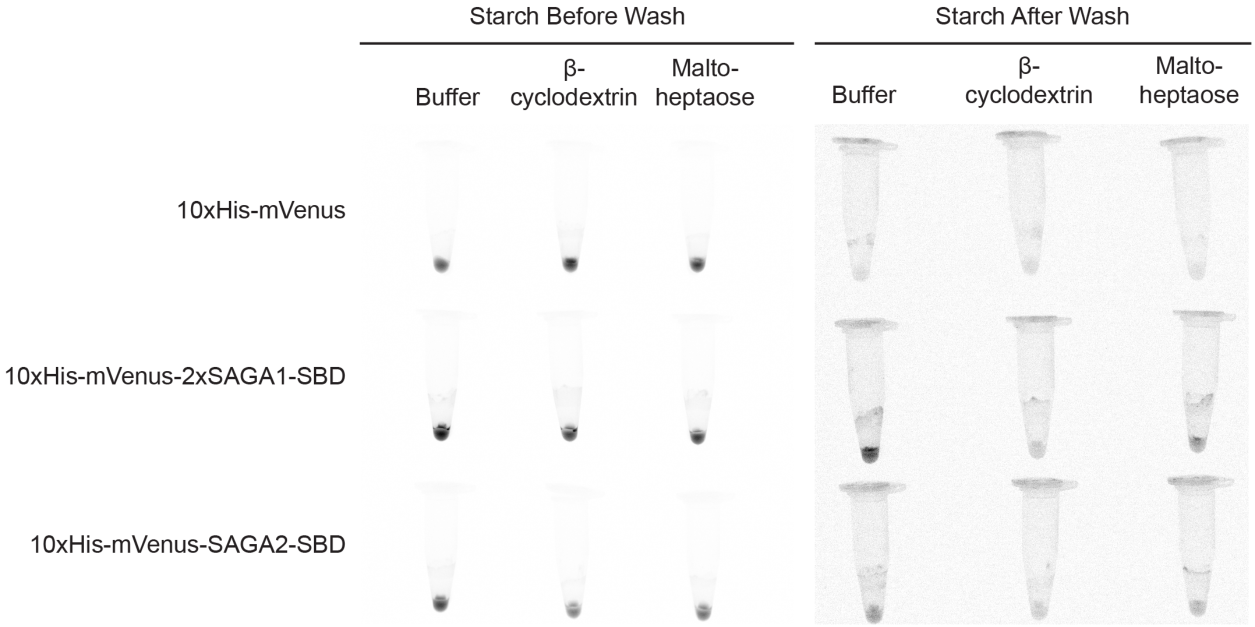

**Figure S16**. Fluorescence images of 10xHis-mVenus, 10xHis-mVenus-2xSAGA1-SBD, and 10xHis-mVenus-SAGA2-SBD incubated with starch pellets in the presence or absence of the CBM20 ligands β-cyclodextrin and maltoheptaose. Following the incubation of each protein in the presence or absence of each CBM20 ligand, the starch granules were pelleted and washed three times. The starch pellets were imaged in microcentrifuge tubes before the first wash (left) and after the final wash (right). The contrast and brightness were adjusted differently in the before- and after-wash images to facilitate comparison of the samples within each group.

**A**

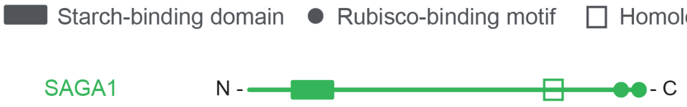

Sorghum bicolor MFP1 XP.02131299.1, Chlamydomonas SAGA1

```
SbMFP1  339   EEKLGEIDVLQEKISLLSQEIDDKAKHIRELSASLSSKEVDYQKLTAFTNETKRSLEL--  396
              EE L EI  LQE +    + I D    + EL+   SS   D Q+ T   +  KRS EL
CrSAGA1 1219  EELLNEIAELQEAVVKYRETISDLNADLAELNQRYSSLVQDRQRAT--EDGGKRSEELGA  1276

SbMFP1  397   ANSRVQQLEEELNTTKNAL  415
              A SR  QL+E+L  +++ L
CrSAGA1 1277  AQSRQAQLQEQLKRSESRL  1295
```

**B**

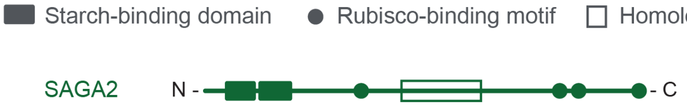

Hordeum vulgare MFP1 KAE8808694.1, Chlamydomonas SAGA2

```
HvMFP1  336   GEMSDLEEKLSLLSQQNDSKEKRIEELNSSLSSKEAEYQN----LRR------FSHQTKE  385
              GE+  ++ +L+  ++ ++S    +  L S+L +  ++YQ     ++R      +S +  E
CrSAGA2 808   GELGLMQSRLAEAARSHNSNSDAVNALESALQAARSQYQGALAEMQRADAAGSYSLRLLE  867

HvMFP1  386   SLEFANA----KIQQLEEEIHTTKNDLASKISSIDSLNEKLQALNSAKKEADEKINELIK  441
               LE A      + + LE+E+   KN  A   S + ++  + Q   S K     ++ E  K
CrSAGA2 868   RLEAAYTGAVDQYEDLEKELVAAKNAQAQSESMLAAM--EAQVATSMKLAGSVQVYE--K  923

HvMFP1  442   ECTDLKASSEMRAN-HDSELLSEKDDLIKQLEEKLSVALSASSKDHEVIAELNKELDSTK  500
              +   L+   +M A  H++EL +        L+ +L+   + +S+  ++   + KE+   K
CrSAGA2 924   QVAVLQEQLQMSAEAHEAELGA--------LQGQLASVEARASQTEDLYESMLKEM---K  972

HvMFP1  501   AMLDDEVAALKSLRDLLKSTEETLSDSRTEVSKLSEDLDEANRMNKDLSLQISNLQSEFN  560
              + LDD +A  + L   L S +    + + +V K +  + E+ R  + L  +++ L+
CrSAGA2 973   SALDDALAREQVLHRELSSMQSRAEEFQRQVEKAAAGMGESTRQEEALRKEVARLRRVIE  1032

HvMFP1  561   EMQEGLTYKLGEAESVCKALSDEVVSAKEMVQKGQEELEATSNGLASAVEARDNLKKELL  620
              + +E L    GE   + ++L  E            E L+A  + L + V  +++   +L
CrSAGA2 1033  QYKEYLEEHDGE---LGRSLERE------------EALDAQLSSLQAVVSQQES---QLR  1074

HvMFP1  621   DVYKKFESTTQELVDERRVVTTLNRELEALAKQLNADSQARKVLEADLDEATRSLDEMNT  680
                 ++ ++ + EL ++RR       +LE   ++L   +  +  +    D  TR +  +
CrSAGA2 1075  GALERQQALSAELEEQRRTADEYQAKLEEQQRELAEAAAKQAAMNERADSMTREMGRLQE  1134

HvMFP1  681   SALSLSKALESTHS  694
                 S    LE++ S
CrSAGA2 1135  LTESYKAKLEASQS  1148
```

**Figure S17**. Sequence alignments of sorghum (*Sorghum bicolor*) and barley (*Hordeum vulgare*) MFP1 homologs with Chlamydomonas SAGA1 (A) and SAGA2 (B). Cartoon depictions of SAGA1 (A) and SAGA2

(B) denote the positions of regions with homology to SbMFP1 (A) and HvMFP1 (B), as well as the CBM20 starch-binding domains and the Rubisco-binding motifs.

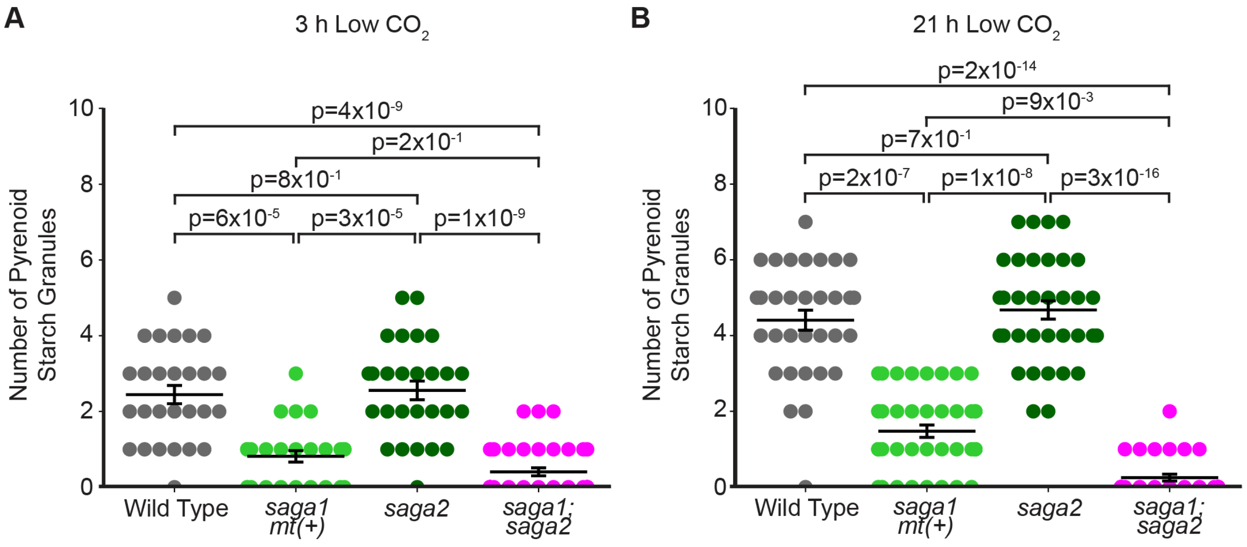

**Figure S18.** Number of starch granules in contact with pyrenoids in wild type, *saga1 mt(+)*, *saga2*, and *saga1;saga2* TEMs (Fig. 3 *B* and *D*, Fig. S8 and S9. (A) Quantifications of the number of pyrenoid starch granules in cells grown at high (3% v/v) $CO_2$ in minimal media, then moved to low (0.04% v/v) $CO_2$ for 3 hours before harvesting. (B) Quantifications of the number of pyrenoid starch granules in cells grown at high (3% v/v) $CO_2$ in minimal media, then moved to low (0.04% v/v) $CO_2$ for 21 hours before harvesting. Statistical p-values were calculated using Kruskal-Wallis ($p=3x10^{-13}$ in A, $p=1x10^{-21}$ in B) followed by Dunn's multiple comparisons test. Error bars represent the standard error of the mean.

**Table S1.** PCR primers used in this study and their sources.

| Primer | Use | Sequence | Source |
|---|---|---|---|
| Fprimer$_{saga1}$ | Verification of the *saga1* insertion | GCATTGAGATCCGAGATGGT | Previously published (1) |
| Rprimer$_{saga1}$ | Verification of the *saga1* insertion | AGTCCAGGCCGACTACTCC | Previously published (1) |
| Fprimer$_{saga2}$ | Verification of the *saga2* insertion | CCACTTGAGAAGCTCTTGGG | Integrated DNA Technologies (IDT) |
| Rprimer$_{saga2}$ | Verification of the *saga2* insertion | GTCCGTGCCTGATCGTAGTC | IDT |
| Rprimer$_{cassette1}$ | Verification of the *saga1* insertion | GCACCAATCATGTCAAGCCT | Previously published (1) |
| Fprimer$_{cassette2}$ | Verification of the *saga2* insertion | CTGCTGAGGCTGAATCTACTC | IDT |
| Rprimer$_{cassette2}$ | Verification of the *saga2* insertion | TCAGTCCTGTAGCTTCATACAAAC | IDT |
| Fprimer$_{minus}$ | Amplification of the *mt(-)* locus | ATGGCCTGTTTCTTAGC | Previously published (2) |
| Rprimer$_{minus}$ | Amplification of the *mt(-)* locus | CTACATGTGTTTCTTGACG | Previously published (2, 3) |
| Fprimer$_{plus}$ | Amplification of the *mt(+)* locus | ATGCCTATCTTTCTCATTCT | Previously published (2, 4) |
| Rprimer$_{plus}$ | Amplification of the *mt(+)* locus | GCAAAATACACGTCTGGAAG | Previously published (2, 4) |

| | | | |
|---|---|---|---|
| Fprimer$_{hyg}$ | Amplification of the APHVII hygromycin resistance gene out of pLM035 | TGCCACCTGACGTCTAAGAA | IDT |
| Rprimer$_{hyg}$ | Amplification of the APHVII hygromycin resistance gene out of pLM035 | TTGTTGAGGCATGCTGAG | IDT |
| Fprimer$_{zeo}$ | Amplification of pLM005-STA2-Venus-3xFLAG for NEBuilder® HiFi DNA Assembly | GGAGAGCAACCCGGGCCCCGT TATGCAGACTCTTGCCGGC | IDT |
| Rprimer$_{zeo}$ | Amplification of pLM005-STA2-Venus-3xFLAG for NEBuilder® HiFi DNA Assembly | GTGGTGGTGCTCGAGGAATT TTACTTGTCGTCATCGTCCT | IDT |

**Table S2.** Sequence alignment BLASTP results for Chlamydomonas SAGA1 and SAGA2 with MFP1 homologs from different cereal species and Arabidopsis. Entries in black text represent BLASTP results that returned at least one alignment between the two proteins. Entries in gray text represent BLASTP results that did not return any significant alignments. Where there were multiple alignments, the E-value, bit score, and percent identity from the best alignment between the two sequences are shown. Note: the homology regions are outside the SAGA1 and SAGA2 predicted CBM20 starch-binding domains.

**Table S3.** Immunoprecipitation-mass spectrometry (IP-MS) dataset. SAGA1-Venus-3xFLAG or SAGA2-Venus-3xFLAG were used as baits. Venus-3xFLAG was used as a control bait to test for non-specific interactions.

**Table S4**. List of Chlamydomonas strains used in this study along with their sources. (*) Strains that tended to clump in liquid culture. (†) The *saga1;saga2;RBCS1-mCherry* strain originally had hygromycin resistance, which was used to select for RBCS1-mCherry expression, but has since lost its ability to grow on hygromycin while continuing to express RBCS1-mCherry, as confirmed using confocal microscopy. (‡) Antibiotics used to select for positive transformants.

| Chlamydomonas Resource Center ID | Strain Description | Mating Type *(mt)* | Source | Antibiotic Resistance |
|---|---|---|---|---|
| CC-4533 | Wild type *CMJ030* | *mt* minus *(-)* | Wild type and parent strain to CLiP library (5) | none |
| CC-5420 | *saga1* | *mt(-)* | Previously published (1) | Paromomycin |
| LMJ.RY0402.088597 | *saga2** | *mt(-)* | CLiP library LMJ.RY0402.088597 (6) | Paromomycin |
| CC-5947 | *saga1 mt(+)** | *mt plus (+)* | Previously published (7) | Paromomycin |
| CC-6341 | *saga1;saga2** | *mt(+)* | Crossing of *saga1 mt(+)* with *saga2* | Paromomycin |
| CC-5422 | *saga1;SAGA1-Venus* | *mt(-)* | Previously published (1, 8) | Paromomycin, Hygromycin |
| CC-6342 | *saga1 mt(+);SAGA1-Venus* | *mt(+)* | Transformation of *saga1 mt(+)* with linearized pRAM118-SAGA1-Venus (1, 7) | Paromomycin, Hygromycin[‡] |
| CC-6343 | *saga1;saga2;SAGA1-Venus** | *mt(+)* | Transformation of *saga1;saga2* with linearized pRAM118-SAGA1-Venus (1, 7) | Paromomycin, Hygromycin[‡] |
| CC-5655 | *CMJ030;SAGA2-Venus* | *mt(-)* | Previously published (8) | Paromomycin |
| CC-6344 | *saga2;SAGA2-Venus** | *mt(-)* | Transformation of *saga2* with | Paromomycin, Hygromycin[‡] |

| | | | | |
|---|---|---|---|---|
| | | | linearized pLM154 (8) | |
| CC-6345 | *saga1;saga2;SAGA2-Venus** | *mt(+)* | Transformation of *saga1;saga2* with linearized pLM154 (8) | Paromomycin, Hygromycin‡ |
| CC-6346 | *CMJ030;SAGA2-Venus;RBCS1-mCherry* | *mt(-)* | Transformation of CMJ030;SAGA2-Venus with linearized pLM035 (9) | Paromomycin, Hygromycin‡ |
| CC-6347 | *saga1;saga2;RBCS1-mCherry;SAGA2-Venus** | *mt(+)* | Transformation of *saga1;saga2;RBCS1-mCherry* with linearized pLM154 (8) | Paromomycin, Hygromycin‡ |
| | *CMJ030;RBCS1-mCherry* | *mt(-)* | Previously published (1, 9) | Hygromycin |
| CC-6348 | *saga1 mt(+);RBCS1-mCherry** | *mt(+)* | Transformation of *saga1 mt(+)* with linearized pLM035 (9) | Paromomycin, Hygromycin‡ |
| CC-6349 | *saga2;RBCS1-mCherry** | *mt(-)* | Transformation of *saga2* with linearized pLM035 (9) | Paromomycin, Hygromycin‡ |
| CC-6350 | *saga1;saga2;RBCS1-mCherry** | *mt(+)* | Transformation of *saga1;saga2* with linearized pLM035 (9) | Paromomycin, Hygromycin†,‡ |
| CC-5374 | *sta6* | *mt(+)* | Previously published (7, 10) | none |
| CSI_FC1G03 | *CMJ030;STA2-Venus* | *mt(-)* | Previously published (11) | Hygromycin |
| CC-6351 | *saga1 mt(+);STA2-Venus** | *mt(+)* | Transformation of *saga1 mt(+)* with linearized pLM006-STA2-Venus-3xFLAG | Paromomycin, Hygromycin‡ |
| CC-6352 | *saga2;STA2-Venus** | *mt(-)* | Transformation of *saga2* with linearized | Paromomycin, Hygromycin‡ |

| | | | | |
|---|---|---|---|---|
| | | | pLM006-STA2-Venus-3xFLAG | |
| CC-6353 | *saga1;saga2;STA2-Venus** | *mt(+)* | Transformation of *saga1;saga2* with linearized pLM006-STA2-Venus-3xFLAG | Paromomycin, Hygromycin[‡] |
| CC-6354 | *CMJ030;STA2-Venus;RBCS1-mCherry* | *mt(-)* | Transformation of CMJ030;STA2-Venus with linearized pLM035 | Paromomycin, Hygromycin[‡] |
| CC-6355 | *saga1;saga2; RBCS1-mCherry; STA2-Venus** | *mt(+)* | Transformation of *saga1;saga2; RBCS1-mCherry* with linearized pChlamy_4-STA2-Venus-3xFLAG | Paromomycin, Hygromycin, Zeocin[‡] |

**Table S5**. Plasmids used in this study and their sources.

| Plasmid | Description/Source | Chlamydomonas Antibiotic Resistance | *E. coli* Antibiotic Resistance | Restriction Enzymes |
|---|---|---|---|---|
| pLM035 (pLM006-RBCS1-mCherry-6xHis) | Rubisco small subunit 1 tagged with mCherry; previously published (9) | Hygromycin | Ampicillin | SpeI |
| pRAM118-SAGA1-Venus-3xFLAG | SAGA1 (Cre11.g467712) tagged with Venus; previously published (1, 7) | Hygromycin | Ampicillin | NdeI |
| pLM154 (pLM099-SAGA2-Venus-3xFLAG) | SAGA2 (Cre09.g394621) tagged with Venus; previously published (8) | Paromomycin | Kanamycin | I-SceI |
| pLM005-STA2-Venus-3xFLAG | STA2 (Cre17.g721500) tagged with Venus; previously published (11) | Paromomycin | Ampicillin | BamHI, MluI |
| pLM006 | Empty mCherry plasmid; previously published (9) | Hygromycin | Ampicillin | BamHI, MluI |
| pLM006-STA2-Venus-3xFLAG | Ligation of double-digested pLM005-STA2-Venus-3xFLAG into double-digested pLM006 | Hygromycin | Ampicillin | EcoRV |
| pChlamy_4 | Thermo Fisher Scientific | Zeocin | Ampicillin | EcoRI |
| pChlamy_4-STA2-Venus-3xFLAG | NEBuilder® HiFi DNA Assembly of PCR-amplified STA2-Venus-3xFLAG into linearized pChlamy_4 | Zeocin | Ampicillin | SspI-HF |
| pGEX-6P-1-SAGA1-2xSBD | NEBuilder® HiFi DNA Assembly of SAGA1-2xSBD into linearized pGEX-6P-1 | N/A | Ampicillin | EcoNI, PspXI |
| pGEX-6P-1-SAGA2-SBD | NEBuilder® HiFi DNA Assembly of SAGA2- | N/A | Ampicillin | EcoNI, PspXI |

| | |
|---|---|
| | SBD into linearized pGEX-6P-1 |

### SI Materials and Methods

#### Measuring Culture Concentrations.

Some strains, noted in Table S4, tended to form clumps in liquid culture. In cases where these clumps impacted the accuracy of the cell counter readings, the cell concentration for each strain being used for the experiment was measured as follows: 1 mL of each culture was transferred to a 1.5 mL microcentrifuge tube. Cultures were centrifuged at 1,000 x g for 5 minutes at RT. The supernatants were discarded and pellets were resuspended in 200 µL autolysin (12) and incubated at RT for 3 minutes to break up cell clumps. Tubes were flicked and inverted ~1x per minute to prevent cells from settling. The cells were then centrifuged at 1,000 x g for 5 minutes and the supernatants were discarded. The pellets were then resuspended in 1 mL of TAP or TP and the cell counts from these resuspensions were measured with a Countess II Automated Cell Counter and used to approximate the cell count of the entire culture.

#### Cloning.

Plasmids used in this study are listed in Table S5. To generate a STA2-Venus-3xFLAG-containing construct conferring hygromycin resistance, pLM005-STA2-Venus-3xFLAG (11) and pLM006 were double-digested with BamHI and MluI (New England Biolabs). The resulting fragments were gel purified with a Macherey-Nagel NucleoSpin Gel and PCR Clean-Up Mini kit and the appropriate fragments were assembled with a New England Biolabs Quick Ligation Kit. To generate a STA2-Venus-3xFLAG-containing construct conferring Zeocin resistance, the STA2-Venus-3xFLAG sequence from pLM005-STA2-Venus-3xFLAG (11) was amplified by PCR (Table S1) to generate 20 bp overhangs for NEBuilder® HiFi DNA Assembly with EcoRI-linearized pChlamy_4 (Thermo Fisher Scientific). Constructs were validated using nanopore sequencing (Plasmidsaurus). To generate plasmids for the CBM20 domain starch binding assay, the following DNA sequences were codon optimized for expression in *E. coli* (Integrated DNA Technologies software) and then further optimized manually for synthesis as gBlocks (Integrated DNA Technologies): a 10xHis-mVenus sequence followed by a $G_4S$ linker, HRV3C cleavage site sequence, and two additional $G_4S$ linker sequences preceded the predicted coding sequences for the CBM20 domains of SAGA1 and SAGA2 plus an additional native sequence after the respective CBM20 domains (Fig. S15*E*), which coded for stabilizing beta strands predicted by AlphaFold. The SAGA1 CBM20 domain was additionally followed by a $3xG_4S$ linker and another copy of the SAGA1 CBM20 domain (Fig. S15*E*). These gBlocks were assembled (NEBuilder HiFi DNA Assembly) into a gel purified (Zymo Research) EcoNI (NEB) and PspXI (NEB) digested pGEX-6P-1 plasmid (Table S5).

**Transformation of Chlamydomonas.**

All plasmids used for Chlamydomonas transformation were linearized with the restriction enzymes listed in Table S5 using standard digestion reactions (New England Biolabs protocols) except pLM154 (*SAGA2-Venus-3xFLAG*) and pRAM118 (*SAGA1-Venus-3xFLAG*). To increase the concentration of DNA in the digestion reaction, 16 µL of purified plasmid (ZymoResearch Classic ZR Plasmid Miniprep Kit) was digested with 1 µL restriction enzyme (New England Biolabs; Table S5) in 1 µL nuclease-free water and 2 µL rCutSmart Buffer (New England Biolabs).

All transformations were performed as described in Hennacy et al., 2024 with some exceptions as follows. To generate *saga2;RBCS1-mCherry* and *saga1;saga2;RBCS1-mCherry*, cells were washed twice in 5 mL of MAX reagent (GeneArt MAX Efficiency Transformation Reagent for Algae, Invitrogen). For all transformations except to generate *saga2;RBCS1-mCherry* and *saga1;saga2;RBCS1-mCherry*, 5 µL of denatured carrier DNA (MB grade from fish sperm, 10 mg/mL, Roche 11467140001) was combined with the linearized plasmids from the digestion reactions before transformation. All centrifugation steps were carried out at 600 x g for 5 minutes. To generate *CMJ030;STA2-Venus;RBCS1-mCherry*, 115 µL of cells were combined with the digestion reaction and carrier DNA for the transformation. To generate *saga2;RBCS1-mCherry* and *saga1;saga2;RBCS1-mCherry*, a co-transformation with both pRAM118-SAGA1-Venus-3xFLAG and pLM035 (Rubisco-mCherry) was attempted, but the resulting transformants only had Rubisco-mCherry expression as confirmed by confocal microscopy. To generate *saga1 mt(+);Rubisco-mCherry*, *saga1 mt(+);STA2-Venus*, *saga2;STA2-Venus*, and *saga1;saga2;STA2-Venus*, a co-transformation with both pLM035 and pLM006-STA2-Venus-3xFLAG was attempted, but the resulting transformants only expressed either STA2-Venus or Rubisco-mCherry and never both. To generate *saga2;SAGA2-Venus* and *saga1;saga2;Rubisco-mCherry;SAGA2-Venus*, since pLM154 conferred paromomycin resistance and the *saga2* and *saga1;saga2;Rubisco-mCherry* strains (Table S4) were already paromomycin resistant, a co-transformation with the hygromycin resistance gene (*AphVII*) was performed. The *AphVII* gene was amplified out of pLM035 by PCR (Table S1) and gel purified (Qiagen). The purified gene was then combined with the digestion reaction so that there was a 10:1 molar ratio of pLM154:*AphVII* for the transformation.

To screen for positive transformations for *CMJ030;STA2-Venus;RBCS1-mCherry*, colonies were transferred into liquid TP in 96-well plates and were grown with agitation under ~100 μmol photons·$m^{-2}$·$s^{-1}$ light for 1 day before screening for positive transformants. For *saga2;RBCS1-mCherry* and *saga1;saga2;RBCS1-mCherry*, colonies were transferred in a 96-array format on TAP 1.5% agar plates supplemented with hygromycin and carbendazim. For all other transformants, colonies were transferred in a 96-array format onto both TAP 1.5% agar plates supplemented with carbendazim and the appropriate antibiotic (paromomycin or hygromycin) and TP 1.5% agar plates with the addition of 250 mg/L amido black (13), supplemented with the appropriate antibiotic (paromomycin or hygromycin), in the same format. The TP amido black plates were placed at 3% (v/v) $CO_2$ under ~100-200 μmol photons·$m^{-2}$·$s^{-1}$ continuous cool white LED light for 4-10 days until colonies were a sufficient size for imaging with an iBright 1500 Imaging System (Invitrogen). Both Venus and mCherry fluorescence were detected using excitation/emission wavelengths of 515-545 nm/568-617 nm, with exposure times of 30, 60, or 90 seconds. Chlorophyll autofluorescence was detected with an excitation/emission of 515-545 nm/710-730 nm and an exposure time of 1 second. Colonies that appeared positive after imaging with the iBright system were then selected for further screening.

**Confocal Microscopy.**

For the dual-tagged Venus and mCherry strains *CMJ030;STA2-Venus;RBCS1-mCherry*, *saga1;saga2;RBCS1-mCherry;STA2-Venus*, *CMJ030;SAGA2-Venus;RBCS1-mCherry*, and *saga1;saga2;RBCS1-mCherry;SAGA2-Venus*, mCherry and Venus signal were collected using the same settings above. Because the 514 nm and 561 nm lasers use different dichroic mirrors on the A1R microscope used for imaging, there was an alignment offset between these two channels for these experiments. To account for this, chlorophyll autofluorescence was collected using both 561 nm/663-738 nm and 514 nm/601-676 excitation/emission. The chlorophyll signal from the 514 nm channel was manually aligned to match the chlorophyll signal from the 561 nm channel to determine the x and y offsets between the two channels. The STA2-Venus channel was then manually adjusted by the same amount as the chlorophyll 514 channel so that it correctly aligned with the Rubisco-mCherry channel for each image.

**Western Blots.**

Cell suspensions (10 mL) were pelleted at 3,428 x g for 10 minutes at 4°C. Pellets were then resuspended in 300 µL of lysis buffer (5 mM Hepes-KOH pH 7.5, 100 mM dithiothreitol, 100 mM $Na_2CO_3$, 2% SDS, 12% sucrose, and cOmplete protease inhibitor cocktail) and transferred to a 1.5 mL microcentrifuge tube. Samples were then heat-denatured in a thermomixer at 37°C, 750 rpm for 10 minutes. Lysates were clarified at 16,000 x g for 5 minutes at 4°C, then aliquoted, flash-frozen in liquid $N_2$, and stored at -80°C until analysis with SDS-polyacrylamide gel electrophoresis.

For loading onto a tris/glycine gradient gel (10%, Bio-Rad Mini-Protean TGX Precast Gel), 100 µL of each lysate was mixed with 100 µL Laemmli buffer (Bio-Rad) supplemented with DTT (900 µL Laemmli buffer + 100 µL 1M DTT). Samples were then heated at 75°C for 10 minutes before loading onto the gel and running at 120 V for ~ 1 hour. Proteins were then transferred to a 0.45 µm polyvinylidene difluoride membrane (Immobilin-P, MilliporeSigma) using a semi-dry transfer system (Bio-Rad) and transfer buffer (20% v/v ethanol, 25 mM Tris, 192 mM glycine, and 0.05% SDS) at 15 V for 40 minutes.

For immunoblot analysis, membranes were blocked in TBST (tris-buffered saline from Bio-Rad + 0.1% Tween-20 from Sigma) containing 5% non-fat dry milk (LabScientific) for 1 hour at RT on a rocking platform. Incubations with the primary antibody (anti-SAGA1 from Meyer et al., 2020, 1:700 dilution) were performed in TBST containing 2.5% milk for 1 hour at RT. Membranes were washed in TBST 3x for 10 minutes on a rocking platform before incubation with the secondary antibody (Goat anti-rabbit IgG H+L HRP, Invitrogen 31466, 1:3,000 dilution) for 1 hour at RT. Membranes were washed again in TBST 3x for 10 minutes on a rocking platform. Immunoreactive proteins were visualized using enhanced chemiluminescence (WesternBright ECL, Advansta) followed by detection using an iBright 1500 Imaging System (Invitrogen).

**Starch Purification and Quantification.**

After growing 1 L bubbling Chlamydomonas cultures, cells were harvested at 3,428 x g for 40 minutes. Supernatants were discarded and pellets were re-suspended in 5 mL TP media and transferred to pre-weighed 50 mL Falcon tubes. Cells were pelleted at 3,428 x g for 10 minutes, then supernatants were discarded, and the pellets were weighed. Pellets were resuspended 1:10 w/v in TP media, then equal volumes for each strain were transferred to a 5 mL Eppendorf tube. Cells were then lysed by sonicating at 4ºC with the following settings: Time: 4 minutes, Pulse: 06 03, Amp: 60%. The lysate was then clarified at 2,000 x g for 20 min. at 4ºC. Supernatants were discarded and the pellets were rinsed with 5 mL 10 mM Tris-HCl, pH 8.0. The pellets were resuspended in 1 mL 10 mM Tris-HCl, pH 8.0, then loaded on top of 5 mL 100% Percoll (Sigma P7828) and centrifuged at 3,428 x g for 20 minutes. The supernatants were then discarded and starch pellets were washed 2x with 5 mL dI water. The supernatants were discarded and then starch pellets were stored at 4ºC overnight. To quantify the extracted starch, pellets were boiled in 25 mL dI water for 3 minutes, then autoclaved for 1 hr at 121ºC to solubilize the starch. The solubilized starch was then quantified following the protocol for the Sigma Starch Assay Kit (SA-20). A Kruskal-Wallis ANOVA test followed by the Conover-Iman multiple comparisons test was used to test for significance in starch coverage between strains and grouped scatter plots were created for data visualization using Python (seaborn.swarmplot).

**CBM20 Domain Protein Purification.**

The plasmids containing the 10xHis-mVenus-CBM20 sequences were expressed in BL21 (DE3) cells (NEB) using an autoinduction system. Cells were grown overnight at 37ºC with shaking at 150 rpm in 3 mL of Miller's LB (Sigma) supplemented with 100 mg/L carbenicillin (GoldBio) and then 1.5 mL of the overnight growth was transferred to 500 mL of LB supplemented with 100 mg/L carbenicillin, 0.05% (w/v) glucose (Sigma), 0.025% (w/v) lactose (Spectrum Chemical), and 10 g of glycerol (Sigma) for growth at 16°C with shaking at 220 rpm. Cells were harvested after approximately 60 hours by centrifugation for 10 minutes at 4,000 x g, resuspended in lysis buffer (50 mM HEPES pH 7.8 [Sigma], 1M KCl [Sigma], 2 mM $MgCl_2$ [Sigma], 5 mM TCEP [GoldBio], 5% [v/v] Glycerol [Sigma], and Roche cOmplete Protease Inhibitor EDTA-free), then frozen at -80°C. Cells were then thawed, lysed on ice using sonication (Qsonica), and the resulting lysate was passed 2x though a 23G needle before centrifugation for 1 hour at 100,000 x g at 4°C to clarify the lysate. Following centrifugation, the supernatant was supplemented with 50 mM imidazole (Thermo Scientific) and filtered using a 0.22 µm vacuum filter (Millipore) before affinity purification using a HisTrap HP column (Cytiva). Following purification, fractions containing the purified proteins were pooled and buffer exchanged into starch binding buffer (50 mM HEPES pH 7.8, 25 mM KCl, 2 mM $MgCl_2$, 5 mM TCEP, 5% [v/v] Glycerol). Protein concentration was determined by using the absorbance at 280 nm and the predicted extinction coefficients for each protein sequence.

**Starch Binding Assay.**

To test whether the SAGA1 and SAGA2 CBM20 domains bind starch, 20 mg of waxy maize starch (Sigma) was washed in 500 µL of starch binding buffer (50 mM HEPES pH 7.8, 25 mM KCl, 2 mM $MgCl_2$, 5 mM TCEP, 5% [v/v] glycerol). The starch was pelleted at 5,000 x g for 30 seconds, the supernatant removed, and then the starch pellet blocked with 100 µL of 5% (w/v) BSA (AG Scientific) for 1 hour with gentle rotation using a tube rotator (VWR). After incubation, the starch was pelleted as before, the supernatant was removed, and the starch pellet was washed 3x with 100 µL of starch binding buffer with the starch pelleted between washes. Following this, 100 µL of each assayed protein at a concentration of 10 µM in starch binding buffer or the same buffer supplemented with 5 mM β-cyclodextrin (Sigma) or maltoheptaose (Sigma) was incubated with the starch for 1 hour with gentle rotation, washed 3x as before, and after the final wash, 100 µL of 1x Laemmli buffer (Bio-Rad) was added to the starch pellet and incubated at 37°C with agitation for 15 minutes. Samples were taken from the initial input protein solution, the supernatant prior to the first wash (unbound protein), the final wash, and from the Laemmli buffer elution. The input and unbound protein samples were diluted 10-fold to prevent overloading on polyacrylamide gels. 10 µL of each sample was loaded onto a Mini-PROTEAN TGX Precast polyacrylamide gel (Bio-Rad) and subsequently stained with InstantBlue Coomassie Protein Stain (Abcam) overnight and destained in water prior to analysis using a 5 second visible light exposure (Invitrogen iBright 1500). To detect the mVenus fluorescence emitted from proteins bound to the starch pellets, the starch pellets before the first wash and after the final wash were imaged on an Invitrogen iBright 1500 using the fluorescent blot setting. The proteins were excited using a 515-545nm filter, and the emission was collected using a 568-617nm filter with an exposure time of 10 msec.

### SI References

1. A. K. Itakura, *et al.*, A Rubisco-binding protein is required for normal pyrenoid number and starch sheath morphology in *Chlamydomonas reinhardtii*. *Proc. Natl. Acad. Sci. U.S.A.* **116**, 18445–18454 (2019).

2. W. Yang, *et al.*, Critical role of *Chlamydomonas reinhardtii* ferredoxin-5 in maintaining membrane structure and dark metabolism. *Proc. Natl. Acad. Sci. U.S.A.* **112**, 14978–14983 (2015).

3. H. Lin, U. W. Goodenough, Gametogenesis in the *Chlamydomonas reinhardtii minus* Mating Type Is Controlled by Two Genes, *MID* and *MTD1*. *Genetics* **176**, 913–925 (2007).

4. J. G. Umen, U. W. Goodenough, Chloroplast DNA methylation and inheritance in *Chlamydomonas*. *Genes Dev.* **15**, 2585–2597 (2001).

5. R. Zhang, *et al.*, High-Throughput Genotyping of Green Algal Mutants Reveals Random Distribution of Mutagenic Insertion Sites and Endonucleolytic Cleavage of Transforming DNA. *Plant Cell* **26**, 1398–1409 (2014).

6. X. Li, *et al.*, A genome-wide algal mutant library and functional screen identifies genes required for eukaryotic photosynthesis. *Nat Genet* **51**, 627–635 (2019).

7. J. H. Hennacy, *et al.*, SAGA1 and MITH1 produce matrix-traversing membranes in the $CO_2$-fixing pyrenoid. *Nat. Plants* **10**, 2038–2051 (2024).

8. M. T. Meyer, *et al.*, Assembly of the algal $CO_2$ -fixing organelle, the pyrenoid, is guided by a Rubisco-binding motif. *Sci. Adv.* **6**, eabd2408 (2020).

9. L. C. M. Mackinder, *et al.*, A repeat protein links Rubisco to form the eukaryotic carbon-concentrating organelle. *Proc. Natl. Acad. Sci. U.S.A.* **113**, 5958–5963 (2016).

10. C. Zabawinski, *et al.*, Starchless Mutants of *Chlamydomonas reinhardtii* Lack the Small Subunit of a Heterotetrameric ADP-Glucose Pyrophosphorylase. *J Bacteriol* **183**, 1069–1077 (2001).

11. L. C. M. Mackinder, *et al.*, A Spatial Interactome Reveals the Protein Organization of the Algal CO2-Concentrating Mechanism. *Cell* **171**, 133-147.e14 (2017).

12. C. D. Amundsen, Autolysin Prep & Assay: Preparation of Autolysin. Available at: https://www.chlamycollection.org/methods/craigs-chlamy-corner/autolysin-prep-assay/.

13. S. Gutiérrez, G. B. Wellman, K. J. Lauersen, Teaching an old 'doc' new tricks for algal biotechnology: Strategic filter use enables multi-scale fluorescent protein signal detection. *Front. Bioeng. Biotechnol.* **10**, 979607 (2022).
